# Fluorogenic Interacting Protein Stabilization for Orthogonal RNA Imaging

**DOI:** 10.1101/2025.01.17.633511

**Authors:** Wen-Jing Zhou, Mei-Yan Wu, Xin-Juan Shao, Li-Juan Tang, Fenglin Wang, Jian-Hui Jiang

## Abstract

Live imaging of RNAs is crucial to interrogate their cellular functions, and approaches allowing fluorogenic RNA imaging with high orthogonality and contrast remains underexplored. We propose a fluorogenic interacting protein stabilization (FLIPS) strategy that enables orthogonal RNA imaging in live cells with fluorescence-labeled RNA interacting proteins (RIPs) stabilized by cognate RNA motifs. We show that the RNA motif stabilizes the otherwise degraded cognate RIP fused to a poly(arginine)- extended C-terminal degron, and the stabilizing effect is enhanced with circular permutation of the RIP. This design affords a generally applicable strategy for different RNA motifs and RIPs, enabling orthogonal and multi-color fluorescence-activated RNA imaging. Our strategy is broadly demonstrated for multiplexed, high-contrast RNA detection and imaging, single molecule RNA imaging and RNA dynamic translocation tracking in live cells. The versatility of our system highlights its potential for interrogating RNA biology and developing RNA-based imaging tools.

---

Live imaging of RNAs provides valuable insights to elucidate their cellular dynamics and functions, and selective fluorescence labeling of RNAs is crucial for their imaging^1^. Current RNA labeling systems rely on two major strategies: multimeric RNA motifs that recruit fluorescent protein-fused RNA-interacting proteins (RIPs) such as MS2/PP7 coat protein (MCP/PCP)^2,3^ and Cas proteins^4-6^, and fluorogenic RNA aptamers that activate otherwise non-fluorescence dyes^7-9^ or degraded FPs^10,11^.

The MCP/PCP system has been established as a gold standard for RNA imaging and mRNA dynamic tracking even at single-molecule resolution^12,13^. A major limitation of this system is the compromised signal-to-background ratio (SBR) due to the excess of unbound fluorescent proteins^11^. Fluorogenic RNA aptamers have emerged as an attractive modality for RNA imaging, as they exploit conditionally fluorescent dyes to minimize background fluorescence^9^. Fluorogenic aptamers have continuously evolved toward multi-colored fluorescence, improved affinity, and enhanced brightness or photostability^14-18^, with demonstrated potential for single-molecule detection^19-21^. An inherent limitation of this strategy is that the dyes are not genetically encoded and require cellular or *in vivo* delivery, and some dyes are susceptible to nonspecific activation in complex biological matrix. To address the issues, a new fluorogenic aptamer has been designed through RNA-stabilized fluorescent proteins fused to a C-terminal TAT peptide-appended degron^10^. This design is completely genetically encodable, precluding the need of dye delivery and enabling use of multi-colored fluorescent proteins. Nonetheless, it requires overlapping between the RNA-binding peptide and the degron, and its extension into orthogonal systems for multiplexed imaging remains elusive.

Here we develop a new concept of fluorogenic interacting protein stabilization (FLIPS), which enables engineering of common RIPs into orthogonal RNA-stabilized fluorogenic proteins for multiplexed RNA imaging. This concept relies on designing fluorescence protein-fused RIPs incorporated with a poly(arginine)-extended C-terminal degron. The RNA motifs are shown to stabilize the designed RIPs through proximity-enhanced electrostatic interactions with the poly(arginine) region, and the stabilizing effect is augmented with circular permutation of these RIPs. The FLIPS system is demonstrated for multiplexed, high-contrast RNA detection, single molecule RNA imaging and RNA dynamic tracking in live cells. We envision that this system will be broadly useful for RNA imaging and novel synthetic biology applications.

## Results

### Design of fluorogenic interacting protein stabilization (FLIPS) system

To develop orthogonal RNA-stabilized fluorogenic proteins, we sought to engineer common RIPs, such as MCP^2^, L7Ae^22^, Cse3^23^, LicT^24^ and LIN28A^25^, into destabilized variants by appending a C-terminal degron (a minimal motif -RG)^25^. The rationale of our design was that binding of RNA motifs to the RIPs could induce steric hindrance on the degron to shield it from recruiting the degradation machinery (Fig. 1a). Hence, we designed to incorporate a poly(arginine) region between the RIPs and the C-terminal degron. As RNA backbones had inherently negative charges to weakly electrostatically interact with poly(arginine), we hypothesized that strong binding between RIPs and cognate RNA motifs could mediate the proximity-dependent association^27,28^ of RNA backbones to the poly(arginine) region, sterically shielding the degron.

**Fig. 1.**
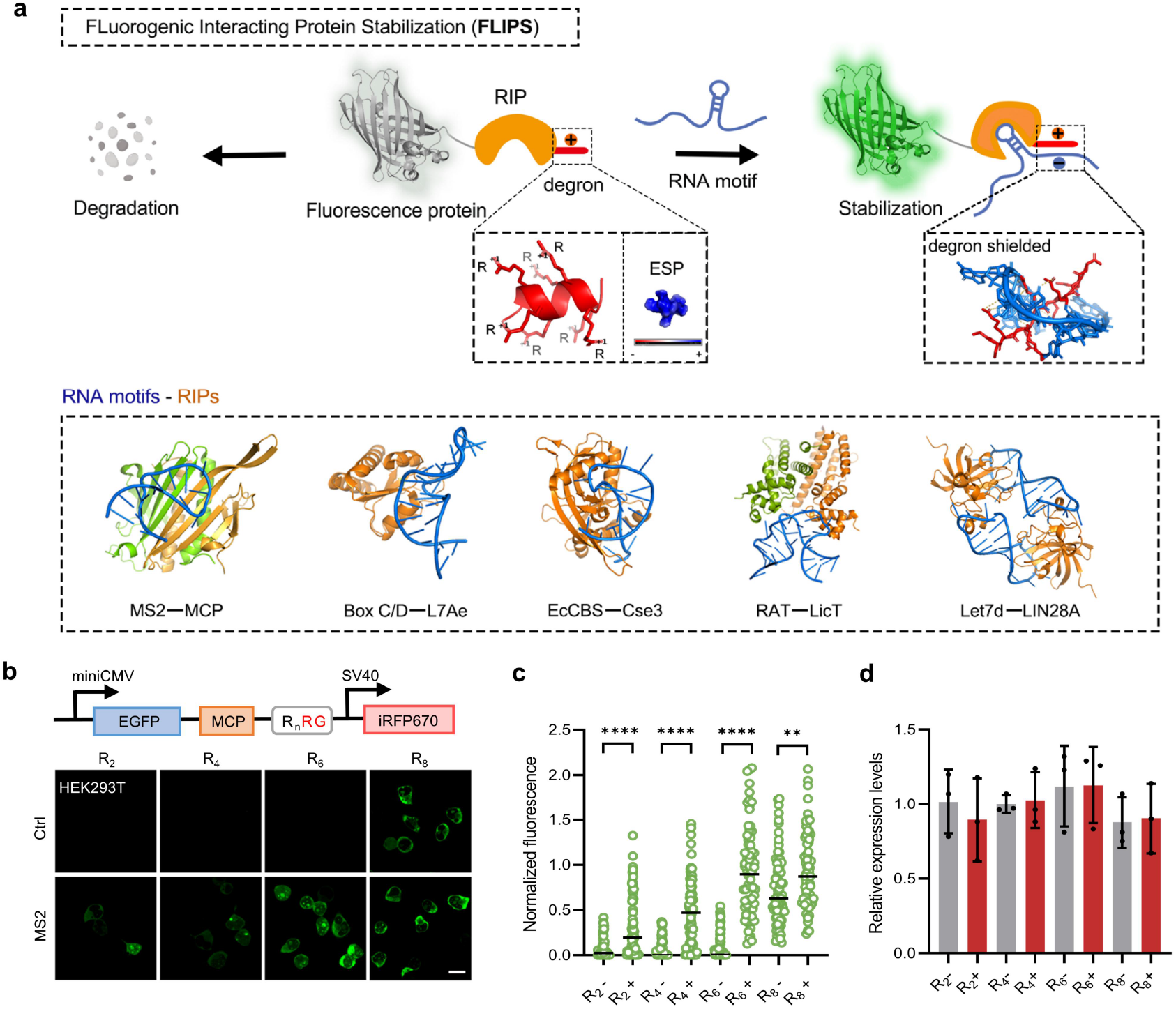
Design and optimization of FLIPS system. **a**, Schematic of the design for FLIPS system. Fluorescence protein fused RIPs tagged with a C-terminal degron are unstable and degraded, whereas binding of cognate RNA motifs shields the degron, stabilizing fluorescent protein fused RIPs with activated fluorescence. A poly(arginine) region (R_n_) is incorporated between the RIP and the degron to increase the interaction between RIPs and RNA motifs. The FLIPS system is generally applicable to different orthogonal RNA motifs and cognate RIPs including MS2−MCP (PDB ID: 2BU1), Box C/D−L7Ae (PDB ID: 1RLG), EcCBS − Cse3 (PDB ID: 4QYZ), RAT − LicT (PDB ID: 6XY4) and Let7d − LIN28A (PDB ID: 5UDZ). ESP: electrostatic potential. **b**, Optimization of arginine numbers in the poly(arginine) region. Confocal images for HEK293T cells co-expressing EGFP − MCP − R_n_RG and circular control RNA (Ctrl, -) or MS2 motif (+). n=2, 4, 6, 8. Scale bar, 10 µm. **c**, Normalized fluorescence intensities for individual cells (100 cells from three independent experiments) in (**b**). Data are normalized to mean fluorescence intensity for cells expressing EGFP−MCP−R_6_RG and circular RNA. Statistical analysis was performed using a two-tailed t-test (**P<0.01, ****P<0.0001). **d**, Relative expression levels of EGFP mRNA determined by qRT-PCR using GAPDH as a control. Data are normalized to mean expression levels for cells expressing EGFP−MCP−R_2_RG and circular RNA. Error bars represent s.d. of three independent experiments.

Motivated by the hypothesis, we engineered the MS2 − MCP based FLIPS system. We designed an EGFP−MCP fusion appended with C-terminal poly(arginine)-extended RG degron (R_n_RG, n=2, 4, 6, 8) through a flexible G_4_S×4 linker. The designed fusions EGFP−MCP−R_2_RG, EGFP−MCP−R_4_RG and EGFP−MCP−R_6_RG displayed the desired destabilized effect with undetectable fluorescence, whereas EGFP−MCP−R_8_RG exhibited unfavorable destabilization (Fig. 1b,c and Supplementary Fig. 1b,c). A proteasome inhibitor experiment showed restored fluorescence for EGFP−MCP−R_2_RG, EGFP−MCP−R_4_RG and EGFP−MCP−R_6_RG (Supplementary Fig. 1d). qRT-PCR confirmed approximate mRNA expressions for these fusion proteins (Fig. 1d). A control with degron-deficient proteins, EGFP−MCP−R_2_, EGFP−MCP−R_4_ and EGFP−MCP−R_6_, validated that C-terminal RG was essential for their degradation (Supplementary Fig. 1e,f). These results revealed that the fusions EGFP−MCP−R_2_RG, EGFP−MCP−R_4_RG and EGFP−MCP−R_6_RG were highly unstable due to a proteasome-mediated degradation. Notably, the undesired destabilization of EGFP−MCP−R_8_RG was presumably ascribed to non-specific interactions of the positive-charge-enriched R_8_RG region with cellular components which sterically shielded the degron.

We then investigated whether the destabilized proteins could be stabilized by MS2 RNA. After co-expression of an MS2-embedded circular RNA^29^, EGFP − MCP − R_2_RG, EGFP − MCP − R_4_RG and EGFP−MCP−R_6_RG delivered fluorescence enhancement by 3.2-fold, 7.4-fold and 12-fold, respectively, relative to a control circular RNA without MS2 (Fig. 1b,c). This result verified that stabilization of these proteins was specific to MS2. A further examination of the linker length between MCP and R_6_RG showed the best SBR of 30-fold achieved with GS linker (Supplementary Fig. 2). A closer look of the crystal structure of MS2 − MCP complex^30^ suggested that a shorter linker yielded closer proximity of the RNA-binding site from the extended degron R_6_RG. To draw the RNA-binding site into closer proximity to the degron, we created MCP circular permutants (R38, A84, A70, D17) by introducing new termini at surface-exposed loops near the binding site and appending the R_6_RG degron at C-termini (Fig. 2a). Based on EGFP fluorescence increases relative to the control RNA, we identified the MCP variant (MCP-A70) stabilized by MS2 with the highest SBR up to 76.2-fold (Fig. 2b,c and Supplementary Fig. 3). We chose this fluorogenic destabilized protein EGFP−cpMCP(A70) −R_6_RG, abbreviated as EGFP−fMCP, for subsequent studies. Additionally, EGFP−fMCP exhibited fluorogenic responses to MS2-embeded circular RNA in different cell lines (Supplementary Fig. 4).

**Fig. 2.**
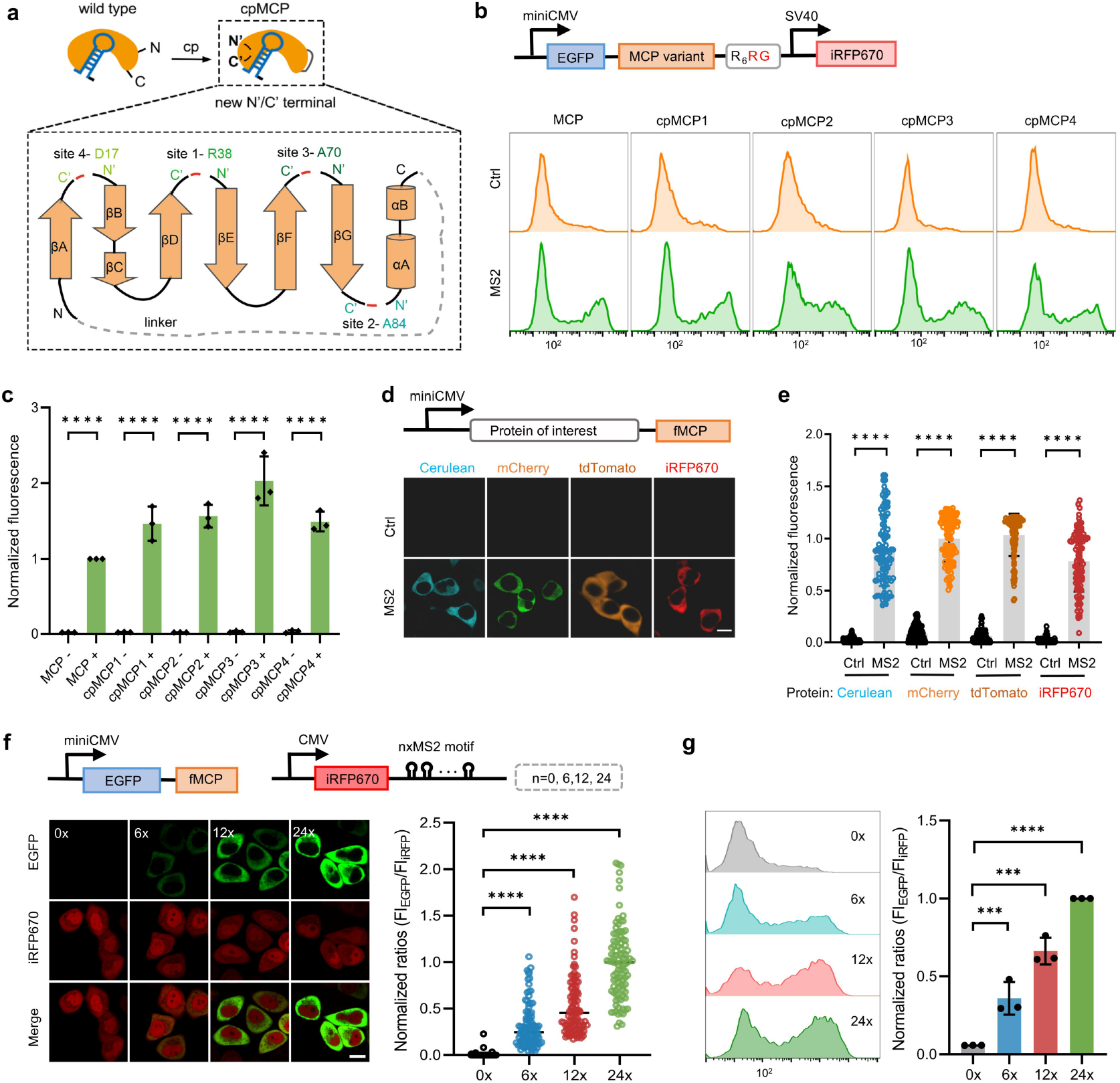
FLIPS system optimization via circular permutation and signal-amplified mRNA imaging. **a**, Schematic showing circular permutation strategy to create fMCP variants with RNA-binding site in close proximity to the degron. The original N and C termini were connected by a linker, and permutation sites were indicated. **b**, Plasmid construct and flow cytometry profiles for HEK293T cells expressing EGFP−cpMCP1-4 −R_6_RG with circular control RNA (Ctrl, -) or circular MS2 (+). **c**, Normalized EGFP fluorescence intensity in (**b**). Data are normalized to mean fluorescence intensity for cells expressing EGFP−MCP−R_6_RG with circular MS2. **d**, Confocal images for cells expressing fMCP fused with different fluorescent proteins for multi-color imaging. **e**, Normalized fluorescence intensities for individual cells (100 cells from three independent experiments) in (**d**). Data are normalized to mean fluorescence intensity for cells expressing Cerulean − fMCP and circular RNA. **f**, Plasmid construct and confocal images for cells co-expressing EGFP−fMCP and MS2-tagged iRFP670 mRNA. Scale bar, 10 µm. **g**, Normalized ratios of fluorescence intensities of EGFP over iRFP670 for individual cells (100 cells from three independent experiments) in (**g**). **h**, Flow cytometry profiles and normalized ratios of fluorescence intensities of EGFP over iRFP for cells expressing EGFP−fMCP and MS2-tagged iRFP670 mRNA. **g**,**h**, Data are normalized to mean ratio for cells expressing EGFP−fMCP and iRFP670−24×MS2. **c**,**e**,**g**,**i**, Statistical analysis was performed using a two-tailed t-test (****P<0.0001, ***P<0.001). Error bars represent s.d. of three independent experiments.

**Fig. 3.**
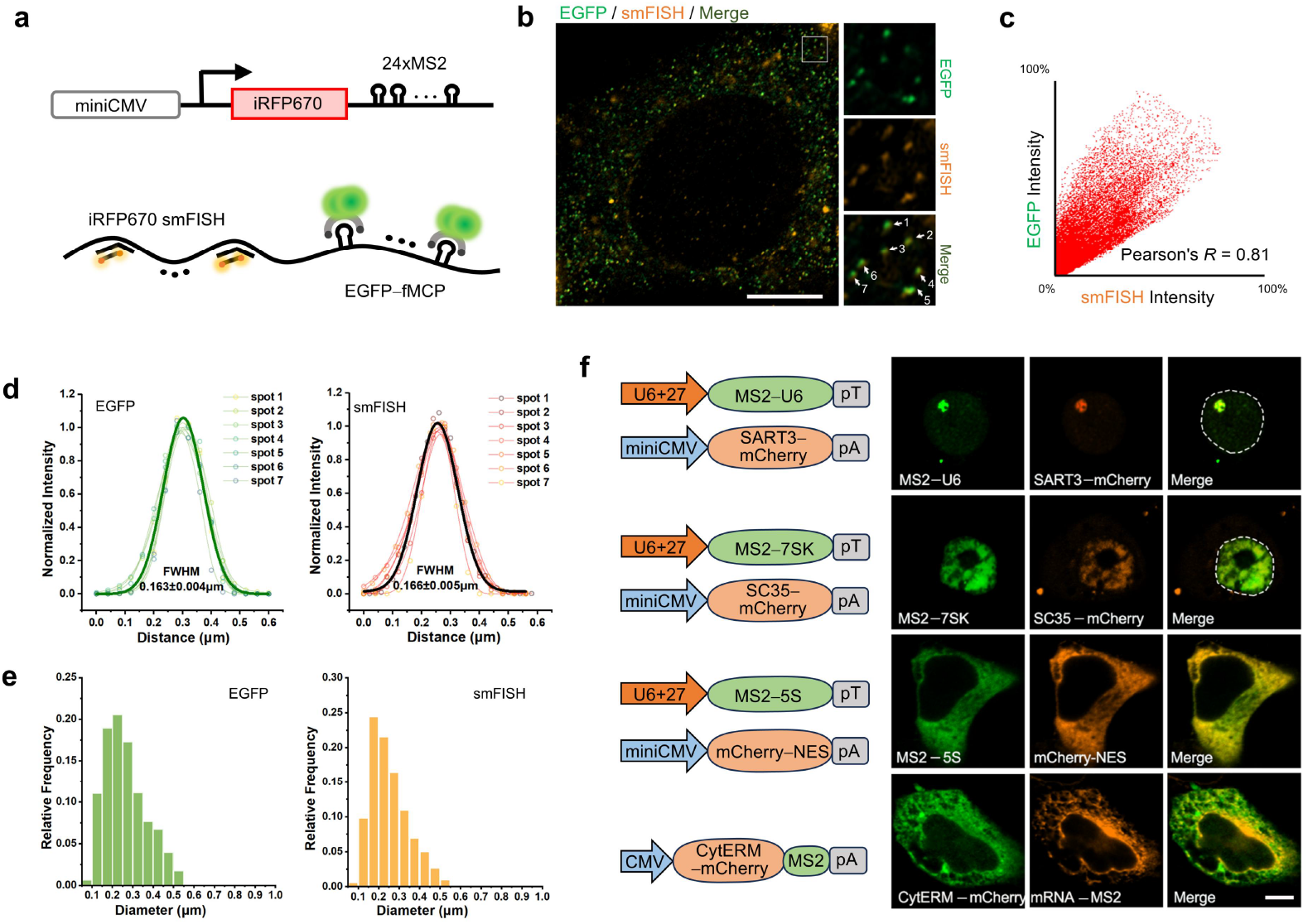
Single-molecule RNA imaging with fMCP system and its effect on subcellular localization of RNAs. **a**, Plasmid construct and schematic of fMCP system for single molecule imaging of iRFP670 mRNA with 24×MS2 tags in comparison to smFISH approach. The transfected MCF-7 cells were hybridized to TAMRA-labeled smFISH probes specific to the iRFP670 sequence. **b**, Confocal images for fixed MCF-7 cells co-expressing fMCP system and colocalization with smFISH probes. Zoom of selected region in the white box were shown on the right. Representative 25 cells from three independent experiments. Scale bar, 5 µm. **c**, Scatter plot showing the co-localization of EGFP and smFISH signals in (**b**). **d**, Normalized fluorescence intensity profiles for EGFP and TAMRA puncta marked by white arrows in (**c**). Symbols, experimental data; thin lines, Gaussian fits to the individual fluorescence intensity profiles; thick line, Gaussian fit to the average profiles, yielding a full width at half maximum (FWHM) of 0.163±0.04 μm for EGFP puncta and 0.166±0.05 μm for TAMRA puncta, respectively. **e**, Histogram of EGFP puncta diameter (Left) and TAMRA puncta diameter (Right), N=1600 puncta in 10 cells from three independent experiments. **f**, Confocal images for MCF-7 cells expressing U6 splicing RNA, 7SK small nuclear RNA, 5S ribosomal RNA or ER-targeting CytERM tagged with MS2 motif and fMCP system. The cells were co-transfected with SART3−mCherry, SC35 −mCherry, or cytosol-localized mCherry to label nuclear speckles, Cajal body and cytosol, respectively. The ER was indicated by CytERM−mCherry. Scale bar, 5 μm. Representative data from three independent experiments.

**Fig. 4.**
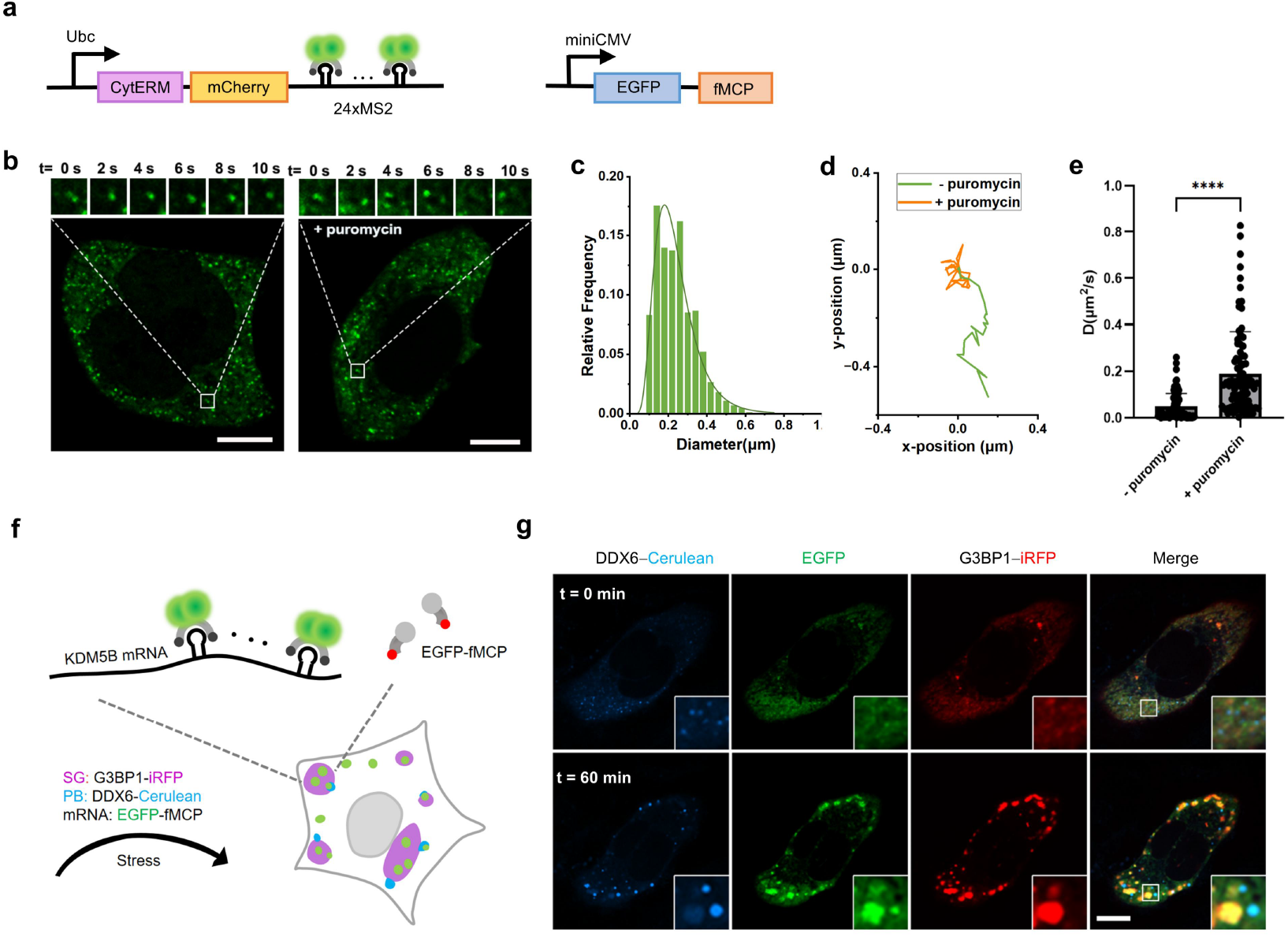
Single-molecule RNA tracking and dynamic imaging of mRNA translocation with fMCP system. **a**, Plasmid constructs for ER-targeting mRNA with 24×MS2 tags and fMCP system. **b**, Confocal images for cells treated with or without a translation inhibitor (100 µg/mL, puromycin). EGFP puncta showed low mobility for transfected cells without puromycin treatment, suggesting that the mRNA was confined to the outer ER membrane. The mobility of EGFP puncta increased for cells treated with puromycin, indicating the mRNA was dissociated from the ER. Scale bar, 5 μm. **c**, Statistical analysis of foci diameter (n = 1600 foci in 10 cells from three independent experiments). **d**, Trajectories of individual RNA tracks for cells treated with or without puromycin at different time points in (**b**). **e**, Analysis of diffusion coefficients of mRNA puncta in cells treated with or without puromycin. Statistical analysis was analyzed using a two-tailed t-test (****P<0.0001). Data are represented as mean ± s.d. (n = 90 puncta from three independent experiments). **f**, Schematic of fMCP system for dynamic imaging of KDM5B mRNA translocation to SG and PB as indicated by G3BP1−iRFP and DDX6−Cerulean, respectively. **g**, Confocal images for transfected cells before and after arsenite (500 µM) induction for 60 min. Zoom of the selected regions in the white boxes were shown on the lower right. Scale bar, 5 μm.

We investigated whether the expressions of other fMCP-fused proteins could also be stabilized by MS2. We obtained consistently undetectable fluorescence in control RNA-expressing cells for fMCP fusions with other fluorescence proteins (Fig. 2d,e) or transcriptional factors (Supplementary Fig. 5). Elevated expressions were obtained for these proteins in MS2-embeded RNA-expressing cells, with 26-to 70-fold enhancement. Together, these results demonstrated that the fMCP system enabled fluorogenic imaging and protein expression control in response to MS2-embedded RNAs.

**Fig. 5.**
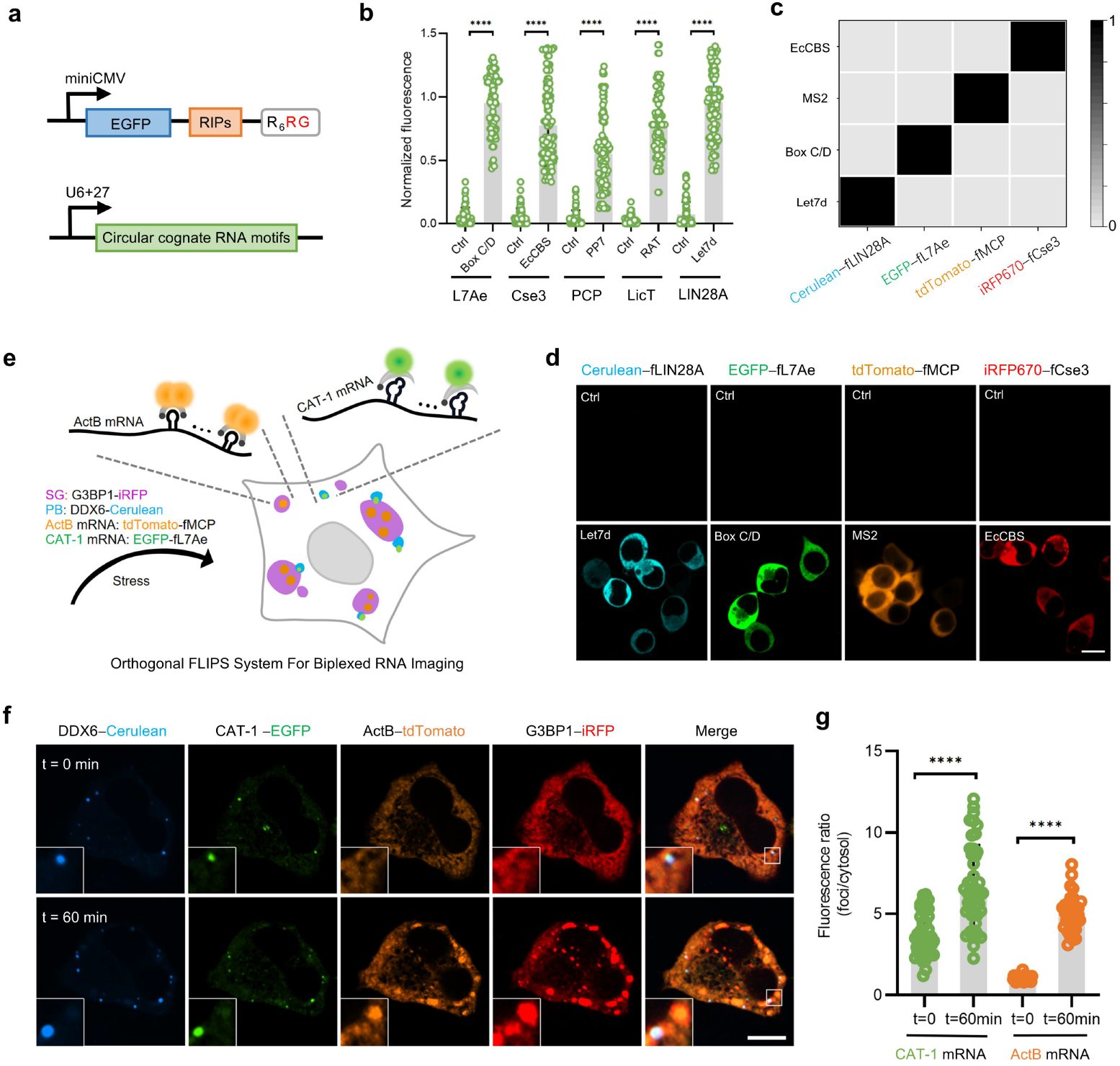
Engineering of orthogonal multi-colored FLIPS system and utility for biplexed RNA imaging. **a**, Plasmid constructs for designing orthogonal FLIPS system using destabilized RIPs and cognate RNA motifs. **b**, Normalized mean fluorescence intensities of HEK293T cells expressing EGFP−RIP−R_6_RG with circular control RNA (Ctrl, -) or circular cognate RNA motif (+). Data are normalized to the mean fluorescence intensity of cells co-expressing EGFP−Lin28A−R_6_RG and circular Let7d. **c**, Heatmap showing orthogonal activation of destabilized RIPs by cognate RNA motifs. **d**, Four-color imaging using FLIPS systems of Cerulean−fLin28A, tdTomato−fMCP, EGFP−fL7Ae and iRFP670−fCse3. Scale bar, 10 μm. **e**, β-actin mRNAs with 8×MS2 motif and CAT-1 mRNAs with 8×Box C/D were tracked by tdTomato−fMCP and EGFP−fL7Ae in MCF-7 cells that expressed the SG marker G3BP1 − iRFP and PB marker DDX6 − Cerulean. **f**, Schematic of orthogonal FLIPS system for simultaneous tracking of β-actin and CAT-1 mRNAs translocation to SG and PB as indicated by G3BP1 − iRFP and DDX6 − Cerulean, respectively. Orthogonal multi-color fluorescence imaging of β-actin mRNA and CAT-1 mRNA before 60 min after arsenite (500 μM) treatment. Zoom of the selected regions in the white boxes were shown on the lower left. Scale bar, 5μm. **g**, Statistical analysis of ratios of EGFP or tdTomato fluorescence intensities in the foci over those in the cytosol (foci/cytosol). Statistical analysis was performed using a two-tailed t-test (****P<0.0001). Data are represented as mean ± s.d.

### mRNA imaging and subcellular localization with fMCP system

We exploited the fMCP system for mRNA imaging using an iRFP670 mRNA target with 0×, 6×, 12× or 24× MS2 in 3’UTR (Fig. 2f). This system generated increased EGFP fluorescence with increasing MS2 motifs, with negligible fluorescence for the control RNA without MS2 (Fig. 2f,g). Besides, the expressions of the MS2-tagged mRNA and iRFP protein exhibited insignificant changes upon EGFP − fMCP co-expression, suggesting that EGFP − fMCP did not alter the stability of MS2-tagged mRNA and its translation (Supplementary Fig. 6). A further investigation using promoters with different expression activities revealed that EGFP fluorescence increased dynamically with iRFP670 − 24×MS2 mRNA expression (Supplementary Fig. 7). These results demonstrated that the fMCP system allowed signal-amplified imaging mRNA in a quantitative manner.

Next, we explored whether the fMCP system enabled single-molecule RNA imaging using low-abundance iRFP670 − 24×MS2 mRNA under miniCMV promoter (Fig. 3a). Distinct EGFP puncta were detected in the cytoplasm, co-localizing with TAMRA puncta obtained using smFISH probes with Pearson correlation coefficient (PCC) 0.81 (Fig. 3b,c). These EGFP and TAMRA puncta showed diffraction-limited size <170 nm and single-mode distribution of fluorescence intensity, indicating that they were single molecules (Fig. 3d,e). Additionally, no EGFP puncta were obtained for the control RNA without MS2 (Supplementary Fig. 8). Besides, distinct fluorescent spots with stronger intensities were observed using a 3×EGFP fused fMCP (Supplementary Fig. 9).

We further compared the fMCP system with a standard PP7−PCP system using low-abundance iRFP−24×MS2−24×PP7 mRNA (Supplementary Fig. 10a). We obtained clear EGFP and mCherry puncta with high co-localization (PCC 0.72), which were verified to be specific to 24×MS2 and 24×PP7 tags (Supplementary Fig. 10b). These puncta exhibited single-molecule characteristic with diffraction-limited size (Supplementary Fig. 10c,d). Notably, EGFP puncta exhibited a higher SBR than mCherry puncta from the PP7−PCP system, and there was no nuclear signal enrichment. Together, these results demonstrated that the fMCP system enabled fluorogenic imaging of single-molecule mRNA with improved contrast.

We investigated whether the fMCP system would affect subcellular localization of MS2-tagged target RNA fusions with varying localization motifs, including U6 splicing RNA^15^, 7SK small nuclear RNA^15^, 5S ribosomal RNA^15^ and ER-targeting CytERM^10^. We observed that U6 splicing RNA fusion co-localized with mCherry-SART3 in Cajal bodies (PCC 0.85), 7SK small nuclear RNA fusion co-localized with mCherry-SC35 in nuclear speckles (PCC 0.92), 5S ribosomal RNA fusion co-localized in the cytoplasm, and ER-targeting mRNA fusion co-localized with CytERM (cytoplasmic end of an endoplasmic reticulum signal – anchor membrane protein) – mCherry on ER membrane (PCC 0.89) (Fig. 3f). Besides, fMCP−tdTomato was specifically recruited to the plasma membrane for cells expressing a target RNA, MS2–Box C/D, tethered to the plasma membrane via L7Ae derived with a CAAX motif (Supplementary Fig. 11). This result testified that our system exhibited negligible effect on subcellular localization of target RNAs.

### Dynamic imaging of mRNA translocation with fMCP system

Our system was further exploited for single-molecule RNA tracking. The 24×MS2-tagged mRNA, tethered via CytERM to ER membrane, were detected as obvious EGFP puncta with diffraction-limited size (Fig. 4a-c). Upon puromycin treatment to dissociate mRNA from ER^10^, EGFP puncta exhibited dramatically increased mobility (Fig. 4b,d). The diffusion coefficients of ER-tethered and dissociated mRNA were estimated to 0.05 and 0.19 µm^2^s^-1^, respectively, according to the mean square displacement (MSD) (Fig. 4e). This result suggested the potential of our system for dynamic tracking of single RNAs.

Next, we demonstrated the fMCP system for dynamic tracking of RNA translocation using KDM5B−8×MS2 mRNA (Fig. 4f), which was reported to translocate into stress granules (SGs) upon induction^31^. Without induction, distinct EGFP puncta were detected in the cytosol (Fig. 4g). Upon arsenite induction, EGFP foci were detected with remarkably increased intensity and size, co-localizing with G3BP1 − iRFP in SGs and DDX6−Cerulean in PBs (Fig. 4g). Time-dependent imaging revealed that SGs formed at 10 min after induction and P bodies grew continuously, accompanied with KDM5B mRNA translocation in SGs and P bodies, and multiple mRNA molecules became continuously accumulated with phase speckles growth over 50 min (Supplementary Fig. 13). This finding was consistent with the literature result for KDM5B mRNA translocation^31^. Additionally, qRT-PCR showed that KDM5B − 8×MS2 mRNA retained almost unchanged expressions under induction (Supplementary Fig. 12b). This result demonstrated the capability of our system for tracking RNA translocation during phase separation.

### Extension of fMCP to orthogonal multi-colored system

Motivated by the fMCP system, we asked whether this design was generally applicable to other RIPs for developing an orthogonal multi-colored fluorogenic RNA imaging system. We chose a set of RIPs with cognate RNA motifs, including L7Ae with Box C/D motif^22^, a CRISPR-associated protein Cse3 with EcCBS (Cse3 binding RNA) motif^23^, PCP with PP7 motif^2^, LicT with RAT motif^24^, and LIN28A with Let7d motif^25^.

We constructed destabilized EGFP-fused RIPs appended with C-terminal R_6_RG degron (Fig. 5a). These destabilized RIPs delivered undetectable fluorescence, but stabilized by cognate RNAs (Supplementary Fig. 14). The SBRs for destabilized L7Ae, Cse3, PCP, LicT and LIN28A were 13-, 11-, 6-, 9- and 10-fold, respectively, testifying that the FLIPS system was feasible for these proteins. We chose the destabilized L7Ae, Cse3 and LIN28A for subsequent optimization, ascribed to their better SBRs.

We turned to circular permutation to further improve SBR for the destabilized RIPs. Five L7Ae circular permutants (H61, K45, P72, V90, D52) were engineered with new termini created near the binding sites (Supplementary Fig. 15a). No circular permutant for L7Ae gave better SBR (Supplementary Fig. 15b,c). Likewise, we circularly permutated five variants of Cse3 (S150, D15, N99, V65, N108), and found the variant Cse3-V65 delivered improved SBR (Supplementary Fig. 16a-c). Meanwhile, we circularly permutated five variants of LIN28A (M17, F50, D88, H98, L105), and found the variant LIN28A-D88 delivered improved SBR (Supplementary Fig. 17a-c). Hence, the improved circular permutants for Cse3 and LIN28A and the wild-type L7Ae were chosen for developing a four-color orthogonal system.

These destabilized RIPs, EGFP − fL7Ae, EGFP − fCse3 and EGFP − fLIN28A, after swapping EGFP for another fluorescence protein such as Cerulean, tdTomato and iRFP670, also exhibited remarkable SBRs (>20) in response to cognate RNAs (Supplementary Fig. 15d,e, 16d,e, 17d,e). Hence, we chose a four-color panel, Cerulean−fLIN28A, EGFP−fL7Ae, tdTomato−fMCP, and iRFP670−fCse3, to investigate their orthogonality. The corresponding fluorescence was found to be specifically activated by cognate RNA motifs, with negligible fluorescence restored by other RNA motifs (Fig. 5c,d). Collectively, these results demonstrated successful construction of the four-color orthogonal FLIPS system for fluorogenic RNA imaging.

### Biplexed RNA imaging with FLIPS system

We demonstrated whether this orthogonal FLIPS system afforded new possibility of biplexed RNA imaging using two destabilized RIPs, tdTomato−fMCP and EGFP−fL7Ae. In parallel to fMCP, we confirmed that the fL7Ae system did not affect subcellular localization of target RNAs (Supplementary Fig. 18), and had the capability for single-molecule RNA imaging (Supplementary Fig. 19).

We then exploited this two-color system to track ActB−8×MS2 and CAT-1−8×BoxC/D mRNAs (Fig. 5e), which were reported to translocate to SGs and PBs, respectively, under stress induction^32,33^. Without induction, rare SGs were detected as indicated by iRFP670 − G3BP1 marker^32^, but clear fluorescence foci of PB marker DDX6−Cerulean^33^ were observed (Fig. 5f). Several CAT−8×BoxC/D mRNA molecules co-localized with PBs.

These observations indicated that without stress induction, ActB and CAT-1 mRNAs were predominantly present in the cytoplasm, with a minority of CAT mRNAs translocated in PBs. After arsenite induction, tdTomato and EGFP foci exhibited increased size, co-localizing with growing SGs and PBs, respectively (Fig. 5f,g). Post-induction time-lapse imaging revealed that remarkable SG and PB growth appeared at 10 min, accompanied with ActB and CAT mRNAs translocated and continuously accumulated in SGs and PBs, respectively, until saturation at 50 min (Supplementary Fig. 21). Additionally, imaging of ActB − 8 × MS2 and CAT − 8 × BoxC/D mRNAs, respectively, using tdTomato − fMCP and EGFP − fL7Ae revealed that individual mRNA exhibited similar translocation behavior (Supplementary Fig. 20). This result was consistent with the previous report. Together, this result demonstrated the ability of our system for biplexed imaging of RNA translocation.

## Discussion

Live cell RNA imaging with multi-color, multiplexing and single-molecule resolution is of persistent challenge. Unlike the proteins, the building blocks of RNAs are non-fluorogenic and could not form fluorophores that are emissive in the visible region. We have developed a concept of RNA-stabilized fluorogenic proteins which allows high contrast, orthogonal and multiplexed RNA imaging in live cells. The system is designed by rendering the fluorescent cognate binding protein unstable via fusing a hybrid degron of positively charged Rs and a degron tag RG. RNA aptamer binding shields the degron, abolishing the degradation of fluorescent cognate protein and delivering activated fluorescence for RNA imaging. We have demonstrated that this design is generally applicable to various RNA motifs and cognate binding proteins. Our system has been successfully employed for multi-color and orthogonal imaging of RNAs, single molecule RNA imaging, simultaneous tracking of RNA translocation dynamics in cells.

Compared to the standard live cell RNA imaging using RNA motifs and fluorescent cognate RNA binding proteins, our system does not have the issue of excessive unbound fluorescent cognate proteins, affording fluorogenic RNA imaging with high contrast. We have demonstrated that our system offers higher S/B ratios as compared to the standard ‘always-on’ systems of using RNA motifs and fluorescent cognate RNA binding proteins. Our system has allowed RNA imaging at single molecule level via tagging the ROIs with 24×RNA motifs. Ascribed to its low background, our system does not need to locate the unbound fluorescent cognate proteins to the nucleus via addition of nuclear localization sequences which allows high-contrast RNA imaging in the cytosol and the nucleus as well.

Compared to the fluorogenic RNA aptamers, our design is completely genetically encodable, allowing RNA imaging without exogenous addition of cognate fluorophores. This complete genetically encodable system is easy to implement and does not have the concerns of non-specific fluorophore activation in complex biological systems. We have repurposed the standard RNA motifs and binding proteins for RNA imaging which have been thoroughly demonstrated for live cell RNA imaging in mammalian cells. Our design is robust and can be easily adapted to RNA imaging without concerns of RNA motif folding efficiency and stability in mammalian cells, which is a formidable challenge for the development of fluorogenic RNA aptamers. Moreover, our system possesses advantages of multi-color, high orthogonality and multiplexing due to the wide availability of RNA motifs, cognate RNA binding proteins and fluorescent proteins. Compared to the recently developed two-color orthogonal RNA imaging using fluorogenic RNA aptamers, we have demonstrated that our design is generally applicable to different RNA motifs and binding proteins, allowing four-color imaging and multiplexed orthogonal RNA imaging in live cells. We have shown that our system enables simultaneous tracking the dynamics of two RNAs to different sub-organelles in live cells.

Overall, as a new RNA imaging approach, our system affords several favorable properties including fluorogenicity, multi-color, orthogonality and multiplexing, highlighting its potential for elucidating RNA biology and developing RNA-based imaging tools.

## Data availability

The datasets generated during and/or analyzed during the current study are available from the corresponding author upon reasonable request. Plasmid DNA and detailed sequence information for described constructs will be made available through AddGene upon acceptance of the publication.

## Acknowledgements

This work was supported by National Key Research Program (2019YFA0905800), NSFC Programs (22090050, 22322404, 22374041, 22274040, 22404050).

## Author contributions

J.H.J. conceived and designed the project. W.J.Z. M.Y.W. and X.J.S. performed the experiments. W.J.Z., L.J.T., F.W. and J.H.J. conducted data analysis and interpretation. J.H.J. and F.W. wrote the manuscript and supervised the project. All authors read and approved the final manuscript.

## Additional information

Supplementary information is available for this paper

**Correspondence and requests for materials** should be addressed to F. Wang and J. H. Jiang.

**Reprints and permissions information is available**.

