## Supplementary material for "Fluorogenic Interacting Protein Stabilization for Orthogonal RNA Imaging"

Table of Contents

I. Materials and Methods ........................................................................................S2

II. Supplementary Tables ........................................................................................S7

III. Supplementary Figures ......................................................................................S10

### I. Materials and Methods

**Materials and reagents.** Opti-MEM, Dulbecco's modified Eagle's medium (DMEM), penicillin, streptomycin, 100% heat-inactivated fetal bovine serum and 4% paraformaldehyde were obtained from Thermo Scientific HyClone (MA, USA). Lipofectamine 3000 was purchased from Xellsamrt (Shanghai, China). MG-132 and puromycin were purchased from Target Mol (Shanghai, China). DEPC-treated $H_2O$, Triton X-100, GeneRuler DNA ladder mix, DNA loading dye, 4S GelRed and agarose were obtained from Sangon Biotech (Shanghai, China). Stellaris hybridization buffer, Stellaris wash buffer A and B were obtained from Dakewe Biotech (Beijing, China). Formamide was purchased form Yeasen Biotechnology (Shanghai, China). Cy3-labeled goat anti-mouse IgG (H+L) and anti-mouse myc antibodies were purchased form Beyotime (Shanghai, China). Arsenite solution was purchased form Leyan Biotechnology (Shanghai, China). RNAex Pro Reagent, RT Mix Kit for qPCR and SYBR Green qPCR Kit were obtained from Accurate Biology (Changsha, China). HeLa cells, MCF-7 cells and HEK 293T were obtained from Cell Bank of Type Culture Collection of Chinese Academy of Sciences (Beijing, China). All the oligonucleotides of HPLC grade (Tables S1 and S3) were synthesized by Sangon Biotech (Shanghai, China). All the other chemicals were of analytical grade and purchased from Sinopharm Chemical Reagent (Shanghai, China). Ultrapure water with an electric resistance >18.25 MΩ was obtained through a Millipore Milli-Q water purification system (Billerica, MA, USA).

**DNA cloning.** PCR Master Mix (Yeasen Biotechnology, 10154ES) was used for PCR amplification. PCR products were separated on 1% agarose gel and the products with correct size were purified with TIANgel Purification Kit (TIANGEN, DP219). Restriction endonucleases were purchased from Takara and used according to the manufacturer's recommended protocol. DNA ligation reactions were carried out using T4 DNA ligase (New England Biolabs, M0202) or Hieff Clone Universal One Step Cloning Kit (Yeasen Biotechnology, 10923ES). DNA plasmids were propagated using chemically competent E. coli DH5α (Sangon Biotech, B528413). Plasmids were purified with TIANpure Mini Plasmid Kit (TIANGEN, DP104) and verified by sequencing in Sangon Biotech.

To construct plasmids for expressing EGFP－MCP fusions with poly(arginine) and RG degron, cDNAs of MCP─$(G_4S)_4$─$R_n$RG (n=2, 4, 6, 8) were obtained by overlap PCR and cloned to the plasmid miniCMV－EGFP－tDeg (Addgene Plasmid #185400) at the BsrGI/ApaI sites to obtain plasmids miniCMV─EGFP─MCP─$R_n$RG (n=2, 4, 6, 8). To construct plasmids for expressing EGFP─MCP─$R_6$RG with different linkers between MCP and $R_n$RG, cDNAs of MCP－GS－$R_6$RG and MCP－$(G_4S)_2$－$R_6$RG were obtained by overlap PCR and cloned to the plasmid miniCMV─EGFP─tDeg at the BsrGI/ApaI sites. To construct plasmids expressing degron-deficient controls, the cDNAs of MCP─$(G_4S)_4$─$R_n$ (n=2, 4, 6, 8) were obtained by overlap PCR and cloned to the BsrGI/ApaI sites. To construct plasmids for expressing EGFP─cpMCP1-4─$R_6$RG, cDNAs for MCP circular permutants with new termini at R38, A84, A70 or D17 were obtained by overlap PCR and cloned to miniCMV─EGFP─tDeg at the BsrGI/ApaI sites. The optimal FLIPS system based on cpMCP3 variant, EGFP－cpMCP3－$R_6$RG, was designated as EGFP－fMCP. Plasmids with iRFP670 as an indicator were obtained by cloning the cDNAs of iRFP670 at the SmaI and BstBI sites under a SV40 promoter. To construct plasmids expressing fMCP fusions with other fluorescent proteins or transcriptional factors, cDNAs of Cerulean, mCherry, tdTomato, iRFP670, Gal4─EGFP, TetR─EGFP and RelA─EGFP were obtained and cloned into the miniCMV─EGFP─fMCP vector at the BsrGI/ApaI sites. Plasmids for expressing FLIPS system using other RIPs were constructed similarly. Briefly, the cDNAs for L7Ae, cpL7Ae1-5, Cse3, cpCse3-1-5, PCP, LicT, LIN28A and cpLIN28A1-5 were inserted into the PCR-linearized miniCMV－EGFP－fMCP plasmid. To construct plasmids for indicating different subcellular localizations, the cDNAs for SART3 － mCherry, SC35－mCherry and mCherry－NES were cloned into the PCR-linearized miniCMV－EGFP－fMCP plasmid, and the cDNAs of G3BP1─iRFP and DDX6─Cerulean were cloned into the plasmid (Addgene #81084) at the NotI and ClaI sites.

To construct plasmids for expressing different circular RNA motifs, the cDNAs for MS2, Box C/D, PP7, EcCBS, Let7d, RAT and a control were obtained and cloned to the Tornado expression plasmid (Addgene plasmid #129405) at the MluI and KnpI sites to obtain vectors U6+27-circular MS2, Box C/D, PP7, EcCBS, Let7d, RAT and control RNA, respectively. The sequences for different RNA motifs were listed in Table S2.

To investigate the ability of the fMCP system for imaging mRNA tagged with MS2 motifs, plasmids CMV－iRFP670－n×MS2 (n=0, 6, 12, 24), cDNAs of n×MS2 (n= 6, 12, 24) were obtained by double digestion from Addgene plasmid #81084 and cloned to plasmid CMV─iRFP670 at the NotI and ClaI sites. Plasmids for expressing CMV－iRFP670－24×MS2 under different promoters including miniCMV, Ubc, SFFV and EF1α were constructed by cloning cDNA of iRFP670 － 24×MS2 to different vectors downstream the corresponding promoters. For constructing plasmids for expressing Ubc─KDM5B─8×MS2, Ubc─ActB─8×MS2, Ubc─CAT-1─8×Box C/D, Ubc─iRFP670─24×MS2─24×PP7, Ubc － iRFP670 － 24×Box C/D, cDNAs of KDM5B － 8×MS2, ActB － 8×MS2, CAT-1 － 8×Box C/D, iRFP670－24×MS2－24×PP7 and iRFP670－24×Box C/D were obtained by overlap PCR and cloned to plasmid (Addgene #81084) at the NotI and ClaI sites.

To construct plasmids for expressing MS2 or Box C/D tagged with different subcellular localization sequences, the cDNAs of MS2－7SK, MS2－U6 and MS2－5S, Box C/D－7SK, Box C/D－U6 or Box C/D─5S were synthesized and cloned into the plasmid (Addgene #129405) at the SalI and XbaI sites. To construct a plasmid for expressing an ER-targeting mRNA reporter, cDNAs for the first 29 amino acids of cytochrome P450, CytERM (MDPVVVLGLCLSCLLLLSLWKQSYGGGKL), and mCherry were cloned into the plasmid (Addgene #81084) at the NotI/NheI sites to obtain plasmid Ubc─CytERM─mCherry─24×MS2.

**Cell culture and transfection.** Cells lines of HeLa, MCF-7 and HEK293T were cultured in DMEM supplemented with 10% fetal bovine serum, 100 U/mL penicillin and 100 g/mL streptomycin at 37°C in a humidified atmosphere containing 5% $CO_2$. The cells were plated on sterilized glass coverslips in 29 mm plates with 14 mm glass bottom well and grown to 30–50% confluency in the corresponding medium at 37°C in a humidified atmosphere containing 5% $CO_2$. The cells were transfected with a mixture of plasmids and Lipofectamine 3000 (2 μL Lipofectamine 3000 per 1 μg plasmids) in Opti-MEM medium according to the manufacturer's instructions. After transfection for 4-6 h, the cells were incubated in a complete medium. Fluorescence images were obtained at a given time or after different treatments.

To optimize the fMCP based FLIPS system, cells were transfected with fluorescent protein-fused MCP with C-terminal degron plasmid (0.1 μg) vectors and circular control (0.2 μg) or MS2 motifs (0.2 μg) plasmids unless otherwise specified. To investigate the ability of fMCP system for imaging mRNAs, cells were transfected with plasmids miniCMV－EGFP－fMCP (0.1 μg) and iRFP670 mRNA (0.2 μg) tagged with different numbers of MS2 motifs driven by different promoters.

To compare the fMCP system and PP7─PCP approach for single-molecule mRNA imaging, MCF-7 cells were transfected with plasmids miniCMV─EGFP─fMCP (0.1 μg), miniCMV─mCherry─PCP─NLS (0.1 μg) and Ubc─iRFP670─24×MS2─24×PP7 (0.2 μg).

To investigate the effect of FLIPS system on the subcellular localization of RNAs, cells were cotransfected with EGFP─fMCP (0.1 μg) or EGFP─fL7Ae (0.1 μg) together with cognate RNA motifs (0.2 μg) tagged with different subcellular localization sequences and a plasmid (0.1 μg) for transfection indication.

To investigate the ability of the fMCP system for dynamic imaging of mRNA translocation, MCF-7 cells were co-transfected with fMCP (0.1 μg), target mRNA (0.2 μg) plasmid and the indicator protein plasmid (0.1 μg).

To extend the FLIPS system to orthogonal multi-colored system, the cells were transfected with EGFP fused RIPs (0.1 μg) and the circular cognate RNA motifs (0.2 μg) plasmids.

To investigate the orthogonal FLIPS system for biplexed RNA imaging, MCF-7 cells were co-transfected with plasmids fMCP (0.1 μg), fL7Ae (0.1 μg), ActB mRNA (0.2 μg), CAT-1 mRNA (0.2 μg) and the indicator protein plasmids (0.1 μg).

**Live cell RNA imaging with the FLIPS system.** Confocal imaging for live cells was performed using a confocal laser scanning fluorescence microscope (Nikon, Ti-E+A1 Si) equipped with a 60× oil objective (NA = 1.49) or 20× air objective (NA= 0.75, low magnification) unless otherwise indicated. For blue channel, samples were excited with a 405 nm laser and signals were collected in the range of 425-475 nm. For green channel, samples were excited with a 488 nm laser and signals were collected in the range of 500-550 nm. For orange channel, samples were excited with a 561 nm laser and signals were collected in the range of 575-625 nm. For red channel, samples were excited with a 640 nm laser and signals were collected in the range of 650-700nm. The images were acquired with a scan speed of 400 Hz, a pixel dwell time of 3.1625 μs and a pinhole size of 26.82 μm using a GaAsP detector. No further data processing was applied unless otherwise indicated.

**Immunofluorescence imaging.** MCF-7 cells were co-transfected with plasmids miniCMV─Gal4/TetR/RelA─EGFP─fMCP and circular control RNA or MS2 motifs for 24 h. The cells were washed with PBS buffer, fixed with 4% paraformaldehyde for 10 min, permeated with 0.2% Triton X-100 for 5 min and blocked with 5% bovine serum albumin for 30 min at room temperature. The cells were incubated with anti-mouse myc antibodies at 37 °C for 60 min. After washing with PBS for three times, the cells were incubated with Cy3-labeled goat anti-mouse IgG at 37 °C for 30 min. Confocal images were obtained by Nikon after washing with PBS for three times. The fluorescence for Cy3 was excited by a 561 nm laser, and the emission signals were collected in the range of 575-625 nm.

**Single-molecule RNA imaging.** To test the ability of the FLIPS system for single-molecule RNA imaging, MCF-7 cells were transfected with miniCMV─iRFP670─24×MS2 and miniCMV─EGFP/3×EGFP─fMCP plasmids, or miniCMV─iRFP670─24×Box C/D and miniCMV─EGFP─fL7Ae plasmids. For comparison with the PP7 ─ PCP system, MCF-7 cells were co-transfected with miniCMV─iRFP670─24×MS2─24×PP7 and miniCMV─EGFP─fMCP with or without co-transfection of mCherry─PCP─NLS. For comparison with smFISH approach, primary smFISH probes comprising a region for hybridization with TAMRA-labeled secondary probes were designed for iRFP670 mRNA. All probes of HPLC purification were obtained from Sangon Biotech (Shanghai, China). Primary probes were pre-hybridized with TAMRA-labeled secondary probes in 1×NEB3 buffer at a final concentration of 40 μM. MCF-7 cells were transfected with miniCMV─iRFP670─24×MS2 or miniCMV─iRFP670 together with miniCMV─EGFP/3×EGFP─fMCP for 24 h. The transfected cells were fixed in 4% paraformaldehyde for 10 min at room temperature followed by permeabilization in 0.2% Triton X-100 for 5 min. The pre-hybridized probes in Stellaris hybridization buffer (100 µL) supplemented with 10% deionized formamide were added to the cells and incubated at 37 °C for 16 h. The cells were washed twice with Stellaris wash buffer A supplemented with 10% formamide and once with Stellaris wash buffer B. Confocal images were acquired with a confocal laser scanning microscope (Zeiss LSM 980) using a 63× oil objective (NA 1.49) and a pixel size of 0.043 µm. Image processing was performed using the ZEN 2 software (Zeiss).

**Single-molecule RNA tracking**. MCF-7 cells were co-transfected with plasmids Ubc—CytERM—mCherry—24×MS2 and miniCMV—EGFP—fMCP. To release the ER-targeting mRNAs, the cells were incubated with a translation inhibitor puromycin (100 µg/mL). Single-molecule tracking was acquired using a spinning disk confocal microscope (Nikon, CSU-W1) with a 100× oil objective lens (NA = 1.49) and a Prime 95B camera. EGFP was excited using a 488-nm laser and emission signals were collected in the range of 500–550-nm. The images were recorded for 30 s with an exposure time of 400 ms (500 ms per frame). The images were acquired with a 1×1 binning, a readout speed of 200 MHz, a disk speed of 4000 rpm and a pinhole size of 50 µm.

**Dynamic Imaging of mRNA translocation**. To investigate the ability of the FLIPS system for imaging mRNA translocation, MCF-7 cells were co-transfected with plasmids Ubc — KDM5B — 8×MS2, miniCMV—EGFP—fMCP, G3BP1—iRFP and DDX6—Cerulean plasmids. After 24 h, the cells were treated with arsenite (500 µM) for 1 h at 37 °C. For tracking the dynamic translocation of KDM5B mRNA under stress, fluorescence images were recorded at an interval of 10 min upon arsenite induction. To investigate the ability of the FLIPS system for biplexed imaging of ActB mRNA and CAT-1 mRNA, MCF-7 cells were co-transfected with plasmids miniCMV — tdTomato — fMCP, Ubc — ActB — 8×MS2, miniCMV—EGFP—fL7Ae, Ubc—CAT-1—8×Box C/D, G3BP1—iRFP and DDX6—Cerulean. After 24 h, the cells were treated with arsenite (500 µM) for 1 h at 37 °C. For tracking the dynamic translocation of ActB mRNA and CAT-1 mRNA under stress, fluorescence images were recorded at an interval of 10 min upon arsenite induction. Fluorescence images were acquired using a spinning disk confocal microscope (Nikon, CSU-W1) with a 100× oil objective lens (NA = 1.49) and a Prime 95B camera. Cerulean was excited with a 405-nm laser excitation and the signal was collected via a 435–485-nm emission filter with an exposure time of 400 ms. EGFP was excited with a 488-nm laser and the signal was collected via a 500–550-nm emission filter with an exposure time of 400 ms. tdTomato was excited using a 561-nm laser and signal was collected via a 580–650-nm emission filter with an exposure time of 400 ms. iRFP was excited using a 640-nm laser and signal was collected via a 665–705-nm emission filter with an exposure time of 400 ms. All the images were acquired with a 1 × 1 binning, a readout speed of 200 MHz, a disk speed of 4000 rpm and a pinhole size of 50 µm.

**Flow cytometry assay.** Cells were detached with 0.25 % trypsin, washed twice with PBS and suspended in PBS. Approximately 1 × $10^4$ live cells from each sample were analyzed using a FACS Celesta Flow Cytometer (BD Biosciences). Fluorescence data were acquired with the following settings: 488 nm laser and 530/30 nm bandpass filter for EGFP, 640 nm laser and 670/30 nm bandpass filter for iRFP670. Data were compensated and analyzed using FlowJo V10 software. Live single cells were gated using forward and side scatter plots (FSC-A versus SSC-A), and forward scatter area and forward scatter height (FSC-A versus FSC-H) to exclude debris and non-singlet events. Flow cytometry data were processed using FlowJo_V10 software.

**Quantitative RT-PCR (qRT-PCR) assay**. The expression levels of EGFP or KDM5B mRNAs were determined by qRT-PCR using GAPDH mRNA as an internal reference. Total RNAs were isolated from HEK293T or MCF-7 cells using RNAex Pro Reagent according to the manufacturer's instructions. RNAs were reverse-transcribed into cDNAs using RT Mix Kit qPCR. The resulting cDNAs were subjected to qRT-PCR assays using SYBR Green qPCR Kit on a BioRad CFX96 system. Primers for target RNAs and GAPDH were designed with the Primer 5 software (Supplementary Table S1). Amplification conditions were one cycle of 95 °C for 30 s followed by 40 cycles of 95 °C for 5 s and 60 °C for 30 s. The specificity of amplification was verified with a final melting-curve analysis step. The data were collected using BioRad CFX manager software. All the samples were normalized to the GAPDH values and the results were expressed as fold changes of cycle threshold (Ct) value relative to the samples without induction using the $2^{-\Delta\Delta Ct}$ method.

**Statistical analysis.** For live cell imaging, all the experiments were performed for at three independent experiments. All data are represented as mean ± s.d. Fluorescence measurements of individual cells were determined using NIS Elements (Nikon) or Fiji software. In single-molecule RNA imaging assays, only puncta with signal at least 1.5-fold brighter than the background signal were analyzed. For KDM5B, ActB or CAT-1 mRNA imaging, foci detection of SGs and PBs were limited to foci at least 2-fold brighter than the background signal. Analysis of fluorescence spot size, brightness and real-time tracking was performed by Fiji software, normalized in Microsoft Excel and plotted in Prism 9.5 (GraphPad) or Origin 2024. Statistical analysis between two groups of samples was evaluated using two-tailed t-tests, with P values ≤0.05 considered statistically significant.

### II. Supplementary Tables

**Table S1. DNA primers for qRT-PCR analysis**

| Name | Sequences (5'-3') |
|---|---|
| EGFP mRNA F primer | CCTCGTGACCACCCTGACCTAC |
| EGFP mRNA R primer | TTGCCGTCGTCCTTGAAGAAGATG |
| GAPDH mRNA F primer | TGGGTGTGAACCATGAGAAGT |
| GAPDH mRNA R primer | TGAGTCCTTCCACGATACCAA |
| KDM5B mRNA F primer | GTGAGCGGTGGGAACGAGTTAA |
| KDM5B mRNA R primer | GGGCAGGGAATGAGTTTCAGCAG |

**Table S2. Sequences for oligonucleotides.**

| Name | Sequences (5'-3') |
|---|---|
| MS2 RNA | ACATGAGGATCACCCATGT |
| Box C/D RNA | GCTCCCGTGATGGCGAAAGCCTGAGGAGC |
| PP7 RNA | TAAGGAGTTTATATGGAAACCCTTA |
| EcCBS RNA | GAGTTCCCCGCGCCAGCGGGGATAAACC |
| Let7d RNA | GATCCGGGGGCTTAGGGCGGGGATTTCGCCCACAAGGAGGTGCCCC CAGATCTA |
| RAT RNA | GGATTGTTACTGCTACGGCAGGCAAAACC |
| U6 | GUGCUCGCUUCGGCAGCACAUAUACUAAAAUUGGAACGAUACAGAG AAGAUUAGCAUGGCCCCUGCGCAAGGAUGACACGCAAAUUCGUGAA GCGUUCCAUAUUUUG |
| 7SK | GGAUGUGAGGCGAUCUGGCUGCGACAUCUGUCACCCCAUUGAUCG CCAGGGUUGAUUCGGCUGAUCUGGCUGGCUAGGCGGGUGUCCCCU UCCUCCCUCACCGCUCCAUGUGCGUCCCUCCCGAAGCUGCGCGCU CGGUCGAAGAGGACGACCAUCCCCGAUAGAGGAGGACCGGUCUUC GGUCAAGG |
| 5S | GUCUACGGCCAUACCACCCUGAACGCGCCCGAUCUCGUCUGAUCUC GGAAGCUAAGCAGGGUCGGGCCUGGUUAGUACUUGGAUGGGAGAC CGCCUGGGAAUACCGGGUGCUGUAGGC |

**Table S3. smFISH probes**

| Name | Sequences (5'-3') |
|---|---|
| secondary probes labeled with TAMRA | TAMRA-CGAGTTCGACGTAGGCATGC-TAMRA |
| iRFP670-sm-1 | GCATGCCTACGTCGAACTCGGAGGTGAGATCGACCTTACG |
| iRFP670-sm-2 | GCATGCCTACGTCGAACTCGTTTTCCGTAATGCGCGTGAT |
| iRFP670-sm-3 | GCATGCCTACGTCGAACTCGAGTTTCGCGTCCAAAGAACG |
| iRFP670-sm-4 | GCATGCCTACGTCGAACTCGAAGTAATCGGCGAGTAGCTC |
| iRFP670-sm-5 | GCATGCCTACGTCGAACTCGCGCTTTGGATCGGAGGACTG |
| iRFP670-sm-6 | GCATGCCTACGTCGAACTCGGTGAGATGTCGAAGGTGCGG |
| iRFP670-sm-7 | GCATGCCTACGTCGAACTCGGATGTACCGTCATGGCGATG |

| | |
|---|---|
| iRFP670-sm-8 | GCATGCCTACGTCGAACTCGCGCAGGCTCGAACTCGATGA |
| iRFP670-sm-9 | GCATGCCTACGTCGAACTCGCGAGCGACTTCAGTTCTTTG |
| iRFP670-sm-10 | GCATGCCTACGTCGAACTCGCGAAGCGGTACAACATCACG |
| iRFP670-sm-11 | GCATGCCTACGTCGAACTCGCGGAAAGTGCTGACCGAGAA |
| iRFP670-sm-12 | GCATGCCTACGTCGAACTCGCGCGTTCTTCAAGTACAGTA |
| iRFP670-sm-13 | GCATGCCTACGTCGAACTCGTGCGCGAACGACAGATCGAG |
| iRFP670-sm-14 | GCATGCCTACGTCGAACTCGCCATGTTCCGCAGAAATTCG |
| iRFP670-sm-15 | GCATGCCTACGTCGAACTCGATGATCGACAGCGACATCGA |
| iRFP670-sm-16 | GCATGCCTACGTCGAACTCGGATGATCAATCCCCATAGCG |
| iRFP670-sm-17 | GCATGCCTACGTCGAACTCGCACGCGGCTCGTAATGATGA |
| iRFP670-sm-18 | GCATGCCTACGTCGAACTCGCGGTGAAGTGCAGCGATAAG |
| iRFP670-sm-19 | GCATGCCTACGTCGAACTCGGATGTGGATCGGCTCGCGAT |
| iRFP670-sm-20 | GCATGCCTACGTCGAACTCGGTCGCAGGCTAGCAGGCAGC |
| iRFP670-sm-21 | GCATGCCTACGTCGAACTCGGTTGCGCAGCGCATGGGCTT |
| iRFP670-sm-22 | GCATGCCTACGTCGAACTCGGTCGCGCCAACCGAAGATCA |
| iRFP670-sm-23 | GCATGCCTACGTCGAACTCGCGTCAGCCGCAGCGGATTGT |
| iRFP670-sm-24 | GCATGCCTACGTCGAACTCGTTCGCCTCGCCGATCACCAT |

Note: The blue base indicates the nucleotides hybridized with the secondary probe.

**Table S4. Amino Acid Sequences for RIPs.**

| Name | Sequences (5'-3') |
|---|---|
| MCP | MASNFTQFVLVDNGGTGDVTVAPSNFANGIAEWISSNSRSQAYKVTCSV RQSSAQNRKYTIKVEVPKGAWRSYLNMELTIPIFATNSDCELIVKAMQGLL KDGNPIPSAIAANSGIY |
| fMCP | AWRSYLNMELTIPIFATNSDCELIVKAMQGLLKDGNPIPSAIAANSGIYGSG MASNFTQFVLVDNGGTGDVTVAPSNFANGIAEWISSNSRSQAYKVTCSV RQSSAQNRKYTIKVEVPKGGSRRRRRRRG |
| L7Ae | MYVRFEVPEDMQNEALSLLEKVRESGKVKKGTNETTKAVERGLAKLVYIA EDVDPPEIVAHLPLLCEEKNVPYIYVKSKNDLGRAVGIEVPCASAAIINEGE LRKELGSLVEKIKGLQK |
| fL7Ae | MYVRFEVPEDMQNEALSLLEKVRESGKVKKGTNETTKAVERGLAKLVYIA EDVDPPEIVAHLPLLCEEKNVPYIYVKSKNDLGRAVGIEVPCASAAIINEGE LRKELGSLVEKIKGLQKGSRRRRRRRG |
| Cse3 | MYLSKVIIARAWSRDLYQLAQGLWHLFPNRPDAARDFLFHVEKRNTPEG CHVLLQSAQMPVSTAVATVIKTKQVEFQLQVGVPLYFRLRANPIKTILDNQ KRLDSKGNIKRCRVPLIKEAEQIAWLQRKLGNAARVEDVHPISERPQYFS GDGKSGKIQTVCFEGVLTINDAPALIDLVQQGIGPAKSMGCGLLSLAPL |
| fCse3 | VATVIKTKQVEFQLQVGVPLYFRLRANPIKTILDNQKRLDSKGNIKRCRVPL IKEAEQIAWLQRKLGNAARVEDVHPISERPQYFSGDGKSGKIQTVCFEGV LTINDAPALIDLVQQGIGPAKSMGCGLLSLAPLGSGMYLSKVIIARAWSRDL YQLAQGLWHLFPNRPDAARDFLFHVEKRNTPEGCHVLLQSAQMPVSTA GSRRRRRRRG |

| | |
|---|---|
| Lin28A | PQLLHGAGICKWFNVRMGFGFLSMTARAGVALDPPVDVFVHQSKLHMEGFRSLKEGEAVEFTFKKSAKGLESIRVTGPGGVFCIGSEDRCYNCGGLDHHAKECKLPPQPKKCHFCQSISHMVASCPLKAQ |
| fLin28A | DRCYNCGGLDHHAKECKLPPQPKKCHFCQSISHMVASCPLKAQGGGGSGGGGSPQLLHGAGICKWFNVRMGFGFLSMTARAGVALDPPVDVFVHQSKLHMEGFRSLKEGEAVEFTFKKSAKGLESIRVTGPGGVFCIGSEGSRRRRRRRG |

Note: The red amino acid sequence indicates the poly(arginine)-extended C-terminal degron.

### III. Supplementary Figures

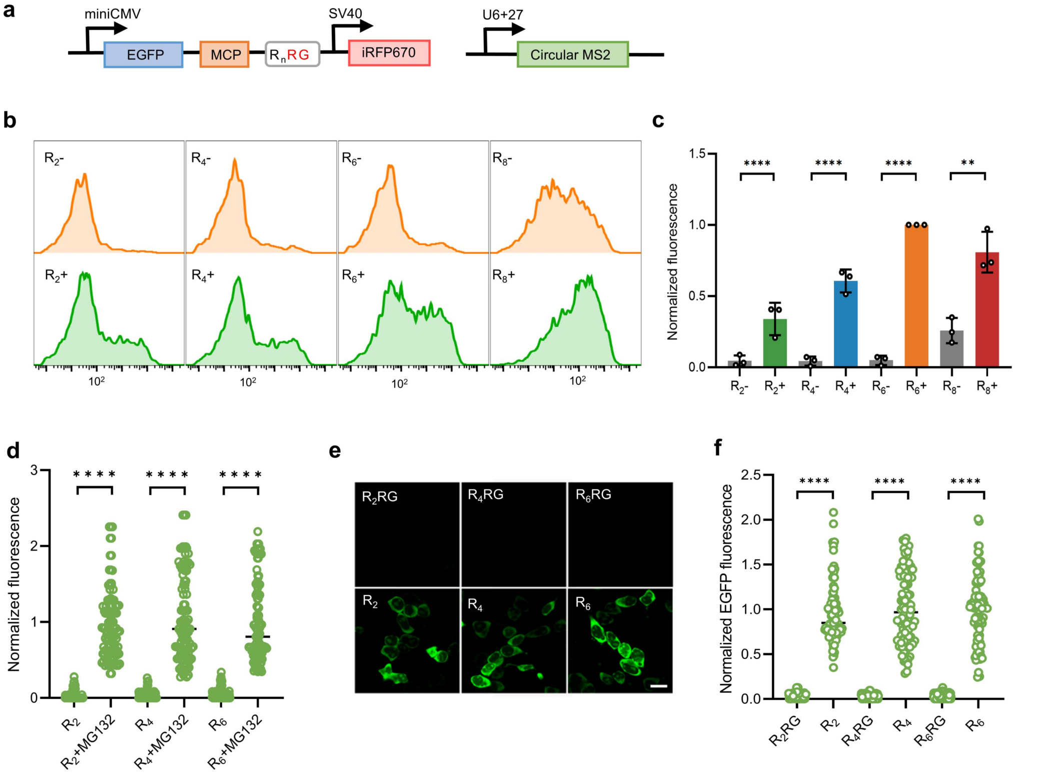


**Supplementary Fig. 1 Optimization of arginine numbers in the poly(arginine) region. a**, Plasmid constructs for expressing EGFP−MCP−$R_n$RG (n=2, 4, 6, 8) and circular MS2. A nuclear export signal was added between EGFP and MCP for cytosol localization. **b**, Flow cytometry profiles for HEK293T cells coexpressing EGFP−MCP−$R_n$RG (n=2, 4, 6, 8) and the circular control RNA or MS2 motif. **c**, Normalized EGFP fluorescence intensities in (**b**). Data are normalized to mean fluorescence intensity for cells expressing EGFP−MCP−$R_6$RG and circular MS2. Statistical analysis was performed using a two-tailed t-test (**P < 0.01, ****P < 0.0001). Error bars represent s.d. of three independent experiments. **d**, Normalized EGFP fluorescence intensities for HEK293T cells expressing EGFP–MCP–$R_n$RG (n=2, 4, 6) with or without MG132 (10 μM) treatment. Data are normalized to cells expressing EGFP–MCP–$R_6$RG treated with MG132. **e**, Confocal images for HEK293T cells expressing EGFP–MCP–$R_n$RG (n=2, 4, 6) or EGFP–MCP–$R_n$ (n=2, 4, 6). Scale bars, 10 μm. **f**, Normalized EGFP fluorescence intensities of individual cells (n= 3 independent cell cultures) in (**e**). Data are normalized to mean fluorescence intensity for cells expressing EGFP–MCP–$R_6$. Data are represented as mean ± s.d from three independent experiments.

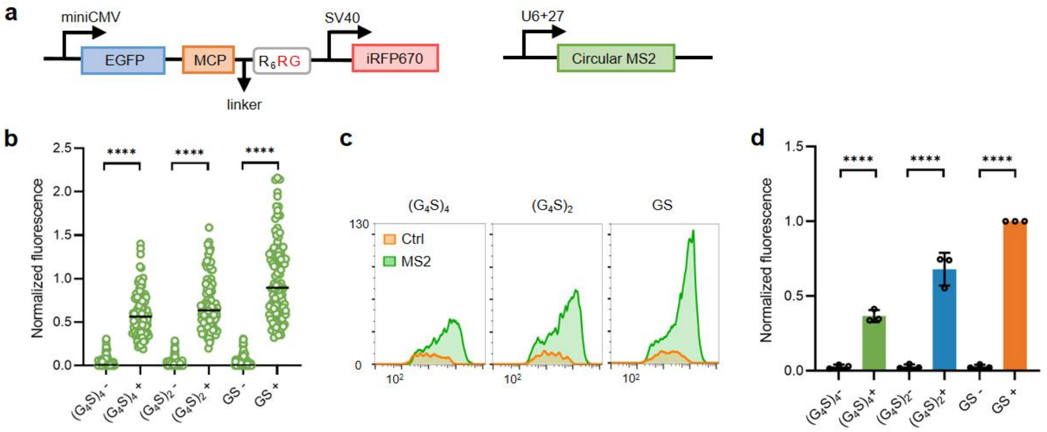


**Supplementary Fig. 2 Linker length optimization between MCP and the poly(arginine) $R_n$ region. a**, Plasmid constructs for expressing EGFP−MCP−$R_6$RG with the linker $(G_4S)_4$, $(G_4S)_2$ or GS between MCP and $R_n$ region. **b**, Normalized fluorescence intensities for cells co-expressing EGFP−MCP−$R_6$RG with different linkers and circular control RNA (Ctrl, -) or circular MS2 motifs (+). Flow cytometry profiles (**c**) and normalized EGFP fluorescence for cells co-expressing EGFP−MCP−$R_6$RG with different linkers and circular control RNA or circular MS2 motifs. **b**,**d**, Data are normalized to cells co-expressing EGFP−MCP−$R_6$RG with the GS linker and circular MS2 motifs. Statistical analysis was performed using a two-tailed t-test (****$P < 0.0001$). Data are represented as mean ± s.d. from three independent experiments.

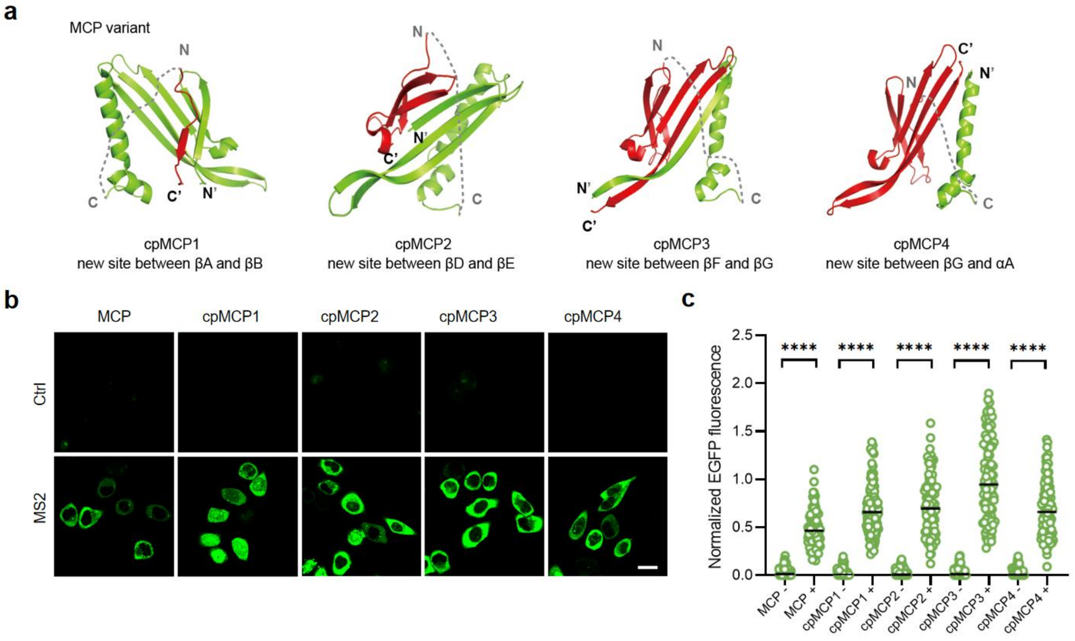


**Supplementary Fig. 3 Optimization of FLIPS system via circular permutation of MCP. a**, Schematic showing different MCP circular permutants with new C terminus as indicated by red. **b**, Confocal images for cells coexpressing EGFP-cpMCP1-4 – $R_6$RG and the circular control RNA (-) or MS2 motif (+). **c**, Normalized EGFP fluorescence of individual cells (n= 100 cells from three independent cell cultures) in (**b**). Data are normalized to mean fluorescence intensity for cells expressing EGFP – MCP – $R_6$R and represented as mean ± s.d from three independent experiments. Statistical analysis was performed using a two-tailed t-test (****$P < 0.0001$).

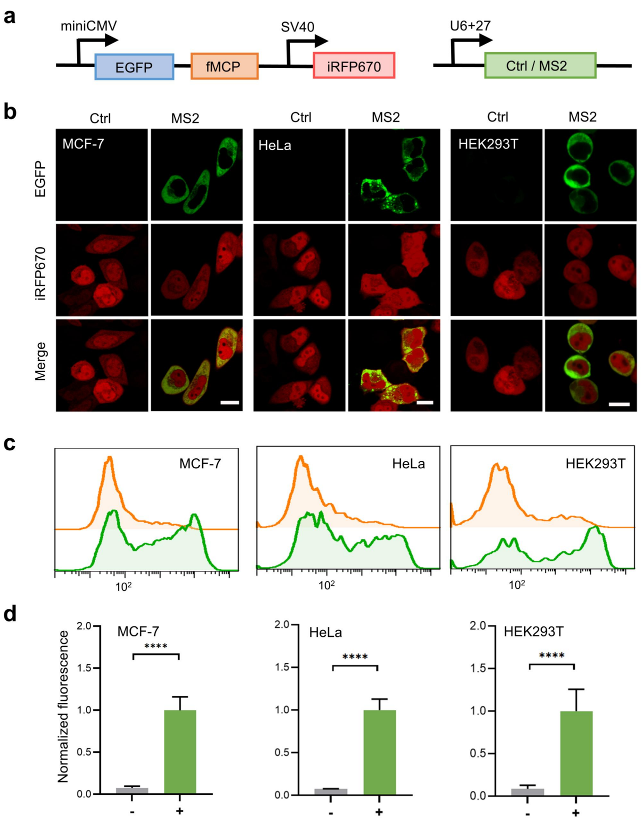


**Supplementary Fig. 4 fMCP system for RNA imaging in different cell lines. a**, Plasmid constructs for EGFP – fMCP system with iRFP for transfection indication. **b**, Confocal images for different cell lines co-expressing EGFP – fMCP and circular control RNA (-) or MS2 motif (+). Scale bars, 10 μm. **c**, Flow cytometry profiles and normalized EGFP fluorescence for different cell lines co-expressing EGFP–fMCP and circular control RNA or MS2 motifs. Data are normalized to each cell line co-transfected with EGFP – fMCP and circular MS2 motifs. Data are represented mean ± s.d. from three independent experiments. Statistical analysis was performed using a two-tailed t-test (****P < 0.0001).

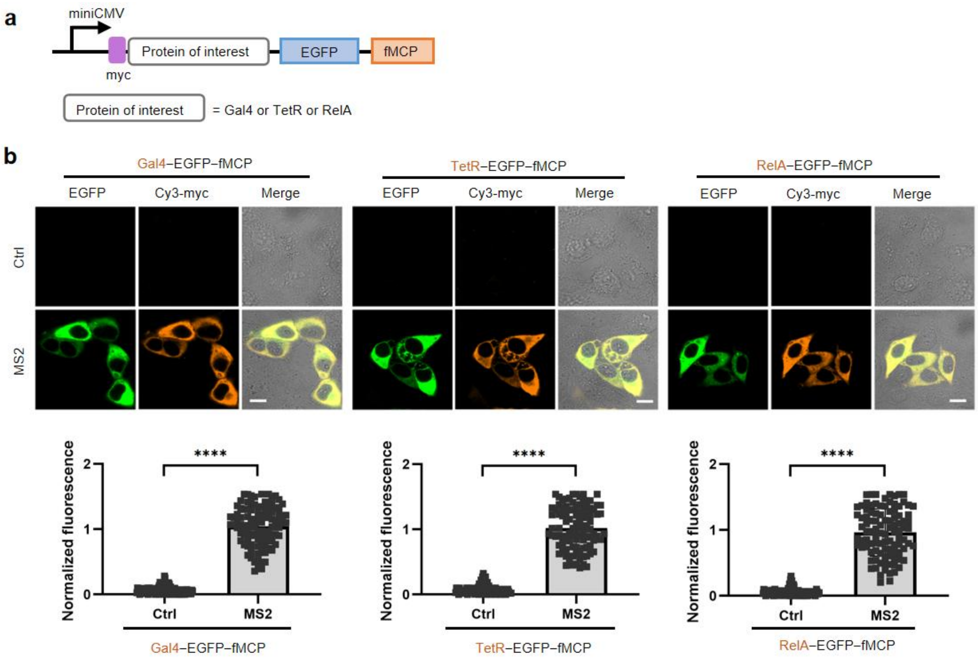


**Supplementary Fig. 5 fMCP system for regulation of different transcriptional factors. a**, Plasmid constructs for expressing EGFP–fMCP fused with myc-tagged transcriptional factors including Gal4, TetR and RelA. **b**, Confocal images for MCF-7 cells co-expressing EGFP – fMCP fused with different transcriptional factors and circular control RNA or MS2 motifs, and immunofluorescence imaging of the myc tag. Scale bars, 10 μm. **c**, Normalized EGFP fluorescence from individual cells (n= 100 cells from three independent experiments) in (**b**). Data are normalized to the mean fluorescence intensities for cells expressing EGFP–fMCP fused with transcriptional factors and circular MS2 motifs. Data are represented mean ± s.d. from three independent experiments. Statistical analysis was performed using a two-tailed t-test (****P < 0.0001).

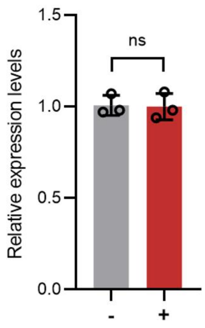


**Supplementary Fig.6 Relative expression levels of iRFP670 mRNA determined by qRT-PCR using GAPDH as a control**. Cells expressed MS2-labeled iRFP670 mRNA alone (-) or co-expressed MS2-labeled iRFP670 mRNA and EGFP−fMCP (+). The result suggested that EGFP−fMCP did not alter the stability of MS2-tagged mRNA and its translation. Data are represented mean ± s.d. from three independent experiments. Statistical analysis was performed using a two-tailed t-test (n.s., not significant).

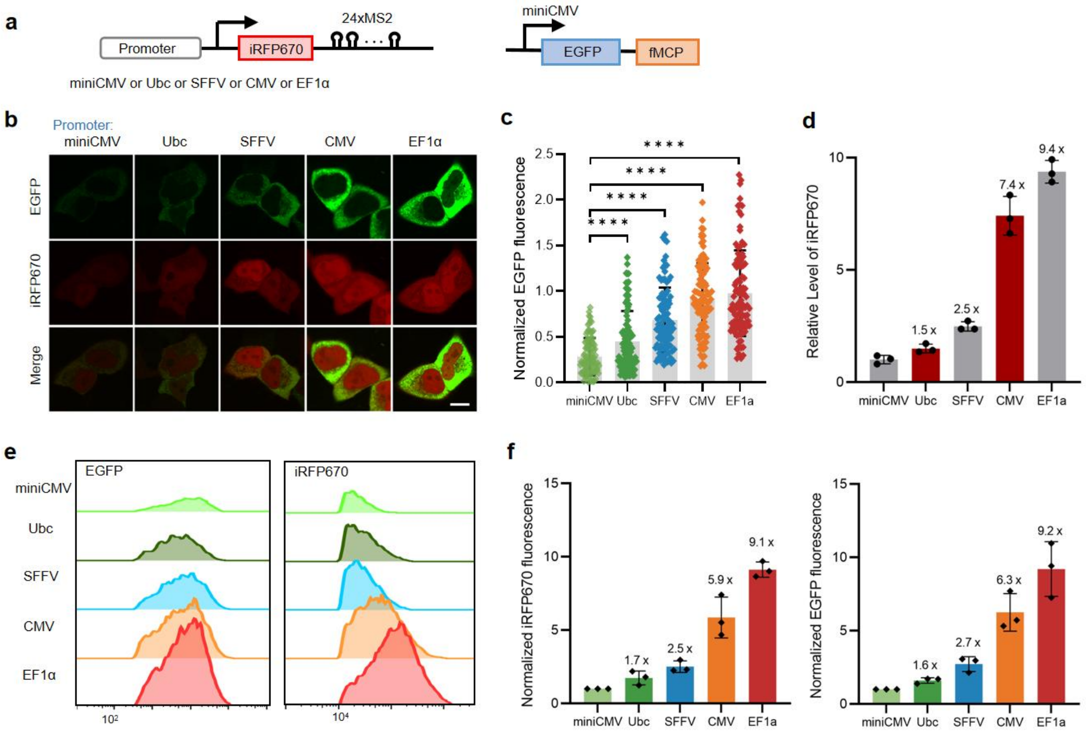


**Supplementary Fig. 7 The fMCP system for imaging mRNAs with different expressions. a**, Plasmid constructs for expressing EGFP – fMCP and iRFP670 – 24×MS2 mRNA under different promoters including miniCMV, Ubc, SFFV, CMV and EF1α. **b**, Confocal images for HEK293T cells coexpressing EGFP–fMCP and iRFP670–24×MS2 mRNA under different promoters. Scale bars, 10 μm. **c**, Normalized EGFP fluorescence from individual cells (n= 100 cells from three independent experiments) in (**b**). **d**, Relative expression levels of iRFP670 mRNA determined by qRT-PCR using GAPDH as a control. Data are represented mean ± s.d. from three independent experiments. **e,** Flow cytometry profiles for cells coexpressing EGFP – fMCP and iRFP670 – 24×MS2 mRNA under different promoters. **f**, Normalized EGFP and iRFP670 fluorescence intensities in (**e**). **c**,**f**, Data are normalized to the mean EGFP or iRFP670 fluorescence intensities for cells expressing EGFP–fMCP and iRFP670–24×MS2 mRNA under miniCMV promoter. Data are represented mean ± s.d. from three independent experiments. Statistical analysis was performed using a two-tailed t-test (****P < 0.0001).

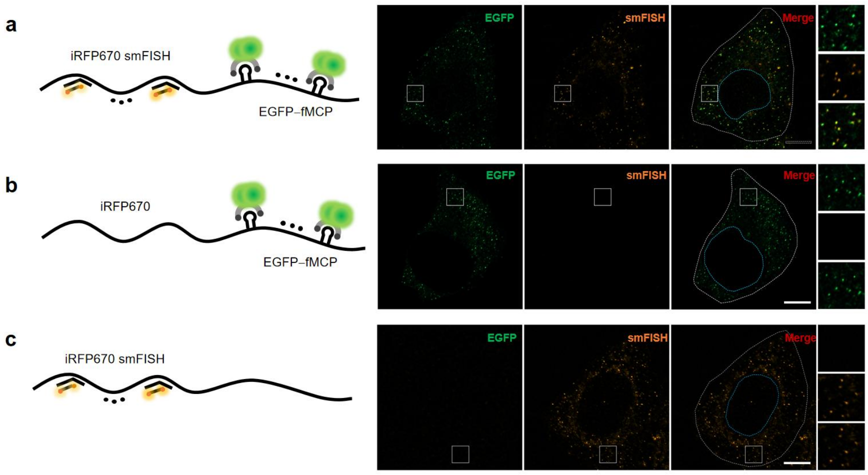


**Supplementary Fig. 8 Specificity of the fMCP system for single-molecule RNA imaging in comparison with smFISH. a**, Confocal images for cells co-transfected with EGFP – fMCP and miniCMV – iRFP670 – 24×MS2, and hybridized with TAMRA-labeled smFISH probes complementary to iRFP670 sequence. Both EGFP puncta from the fMCP system and TAMRA puncta from smFISH probes were observed with good co-localization. **b**, Confocal images for cells expressing EGFP – fMCP and miniCMV – iRFP670 – 24×MS2. Fluorescence labeling diagram and fluorescence imaging of cells coexpressing EGFP – fMCP plasmids, and treated with random TAMRA-labeled smFISH probes not complementary to iRFP670 sequence. Only EGFP puncta from the fMCP system were observed. **c**, Confocal images for cells transfected with EGFP – fMCP and miniCMV – iRFP670, and hybridized with TAMRA-labeled smFISH probes complementary to iRFP670 sequence. Only TAMRA puncta from smFISH probes were observed. **a,b, c,** Zoom of the selected regions in the white boxes were shown on the right. Scale bars, 5 μm. Representative images (n=20 cells) from three independent experiments.

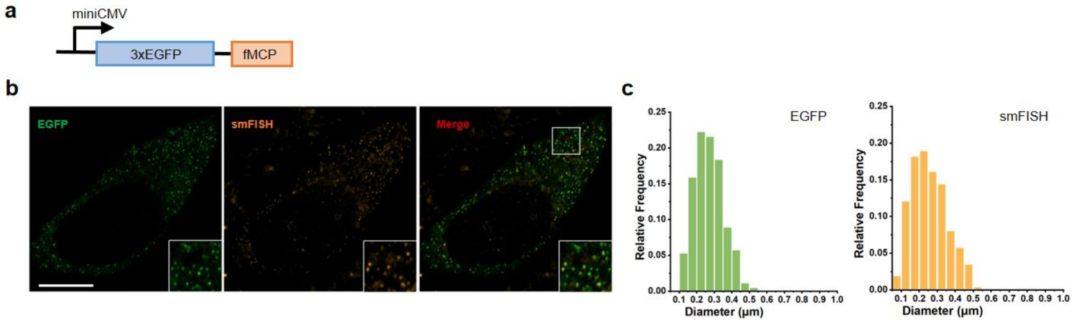


**Supplementary Fig. 9 Single-molecule mRNA imaging using the 3×EGFP – fMCP system in comparison with smFISH.** **a**, Plasmid construct for 3×EGFP–fMCP. **b**, Confocal images for MCF-7 cells co-transfected with plasmids miniCMV – iRFP670 – 24×MS2 and 3×EGFP – fMCP, and hybridized with TAMRA-labeled smFISH probes complementary to iRFP670 sequence. Zoom of the selected regions in the white boxes were shown on the lower right. Scale bars, 5 μm. **c**, Histogram of EGFP puncta diameter (Left) and TAMRA puncta diameter (Right), N=1800 puncta in 10 cells from three independent experiments.

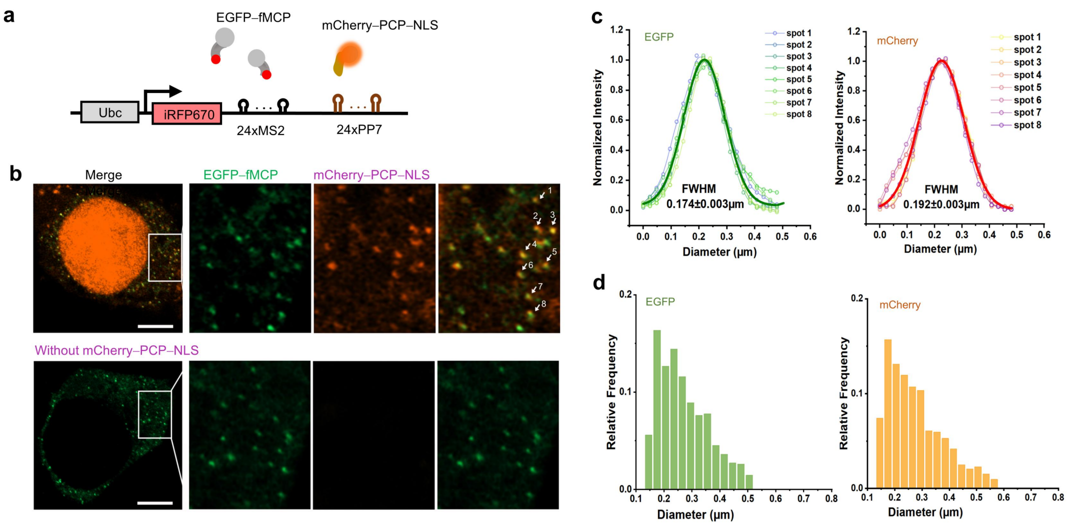


**Supplementary Fig. 10 Comparison of the fMCP system and PP7 – PCP approach for single-molecule RNA imaging. a,** Schematic diagram for imaging iRFP670 mRNA tagged with 24×MS2 and 24×PP7 using the fMCP system and PP7 – PCP approach. **b**, Confocal images for MCF-7 cells expressing iRFP670 – 24×MS2 – 24×PP7 and EGFP – fMCP with or without co-expressing mCherry – PCP – NLS. Scale bar, 5 μm. **c**, Normalized fluorescence intensity profiles for EGFP and mCherry puncta marked by white arrows in (**b**). Symbols, experimental data; thin lines, Gaussian fits to the individual fluorescence intensity profiles; thick line, Gaussian fit to the average profiles, yielding a full width at half maximum (FWHM) of 0.174±0.03 μm for EGFP puncta and 0.192±0.03 μm for mCherry puncta, respectively. **d**, Histogram of EGFP puncta diameter (Left) and mCherry puncta diameter (Right), N=1600 puncta in 10 cells from three independent experiments.

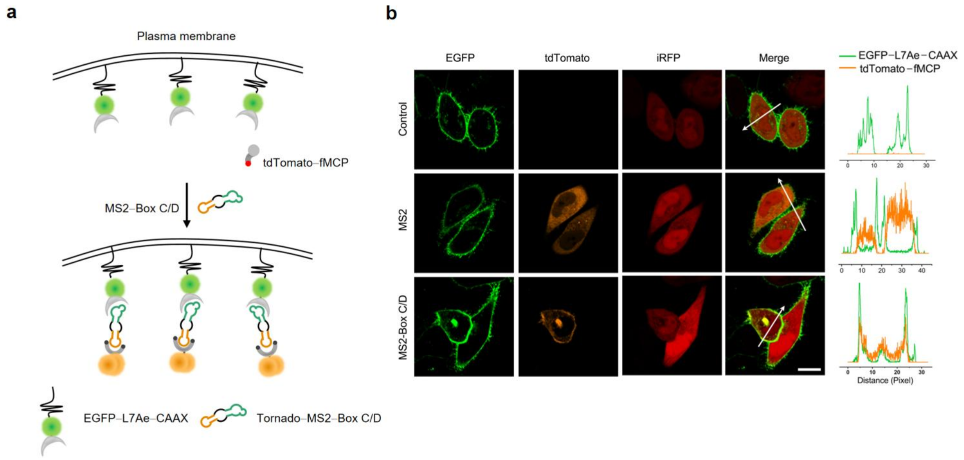


**Supplementary Fig. 11 Specific stabilization of fMCP system at the plasma membrane. a**, Schematic diagram showing the stabilization of tdTomato − fMCP by circular MS2 − Box C/D motifs tethered to the plasma membrane via EGFP−L7Ae fusion with a CAAX motif. **b**, Confocal images and localization distributions for tdTomato in MCF-7 cells co-expressing tdTomato−fMCP, EGFP−L7Ae−CAAX, and circular control RNA, circular MS2 motifs or circular MS2−Box C/D motifs. Fluorescence intensity profiles along the white arrows were shown on the right. Scale bar, 10 μm.

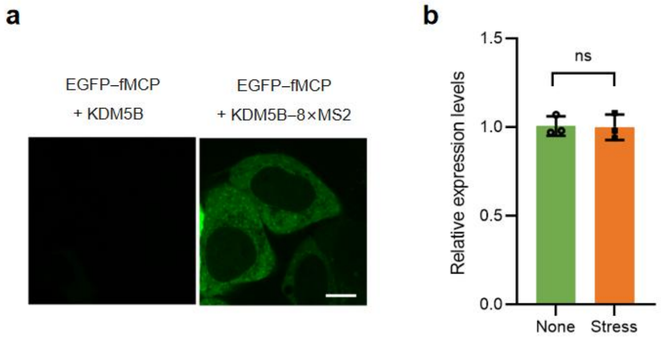


**Supplementary Fig. 12 Dynamic imaging of KDM5B mRNA translocation with fMCP system. a**, Confocal images for MCF-7 cells co-expressing EGFP−fMCP and KDM5B or KDM5B−8×MS2. EGFP fluorescence was specifically restored for cells expressing KDM5B tagged with MS2 motifs. Scale bar, 5 μm. **b**, Relative expression levels of KDM5B mRNA before and after arsenite (500 μM) induction for 60 min as determined by qRT-PCR using GAPDH as a control. Data are represented mean ± s.d. from three independent experiments. Statistical analysis was performed using a two-tailed t-test (n.s., not significant).

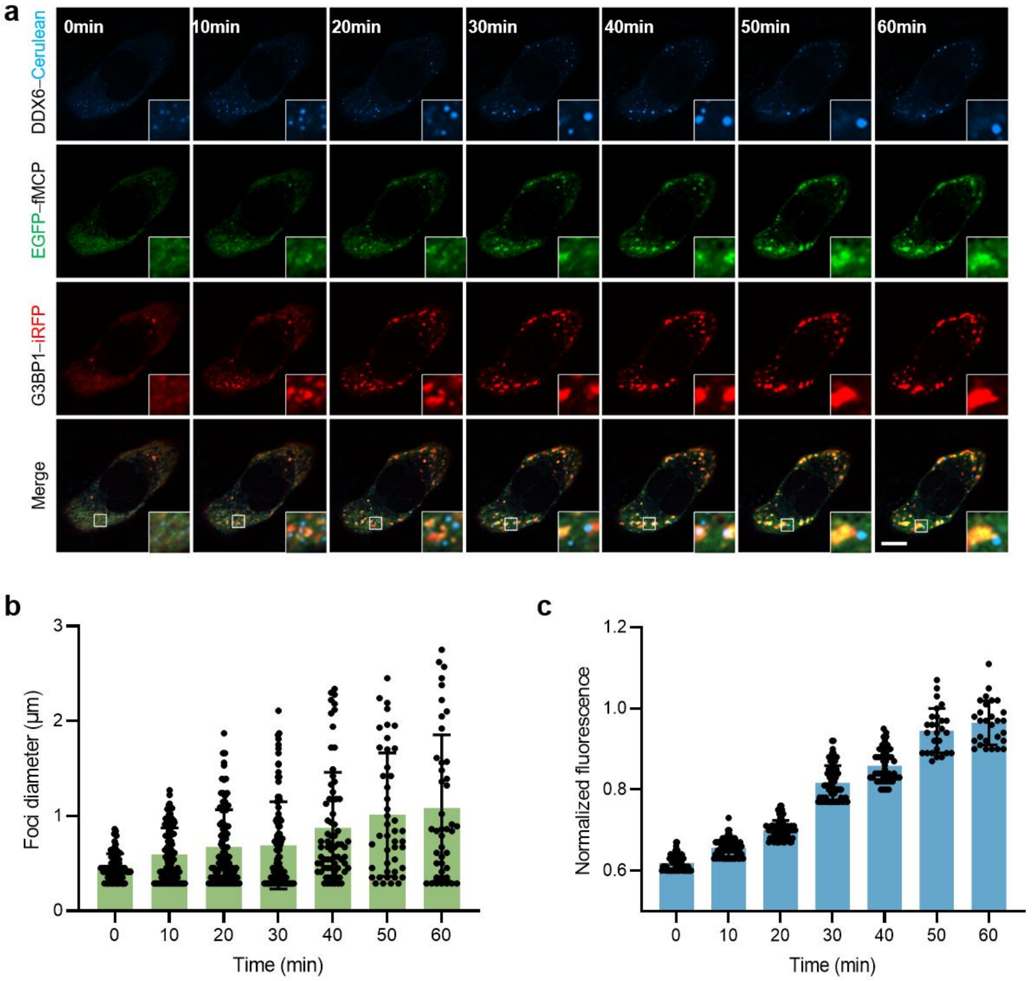


**Supplementary Fig. 13 Real-time imaging of KDM5B mRNA translocation dynamics using fMCP system. a**, Confocal images of cells expressing KDM5B – 8×MS2 and EGFP-fMCP at different time intervals upon arsenite (500 μM) induction. Zoom of selected regions in the white boxes were shown on the lower right. Scale bar, 5 μm. Statistical analysis of foci diameter (**b**) and normalized EGFP fluorescence intensities (**c**) of SGs for cells under arsenite induction. Data are normalized to the EGFP fluorescence intensities at 60 min after arsenite induction and are represented as mean ± s.d. from three independent experiments.

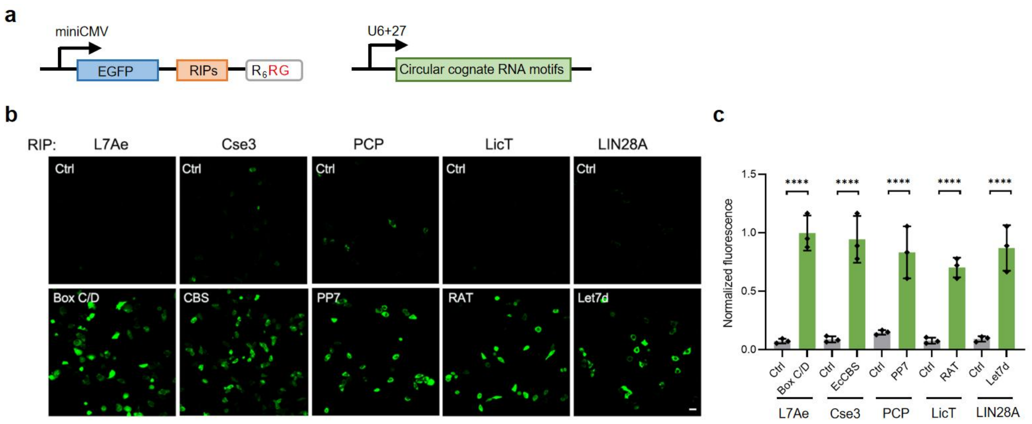


**Supplementary Fig. 14 Orthogonal FLIPS system using other RIPs and cognate RNA motifs. a**, Plasmid constructs for designing orthogonal FLIPS system using destabilized RIPs and cognate RNA motifs. **b**, Confocal images for cells expressing EGFP – RIPs – $R_6RG$ with circular control RNA (-) or circular cognate RNA motifs. Scale bar, 5 μm. **c**, Normalized mean EGFP fluorescence intensities for cells in (**b**) plotted from flow cytometry profiles. Data are normalized to mean EGFP fluorescence intensity for cells expressing EGFP–LIN28A–$R_6RG$ and circular Let7d. Statistical analysis was performed using a two-tailed t-test (****P < 0.0001). Data are represented as mean ± s.d. from three independent experiments.

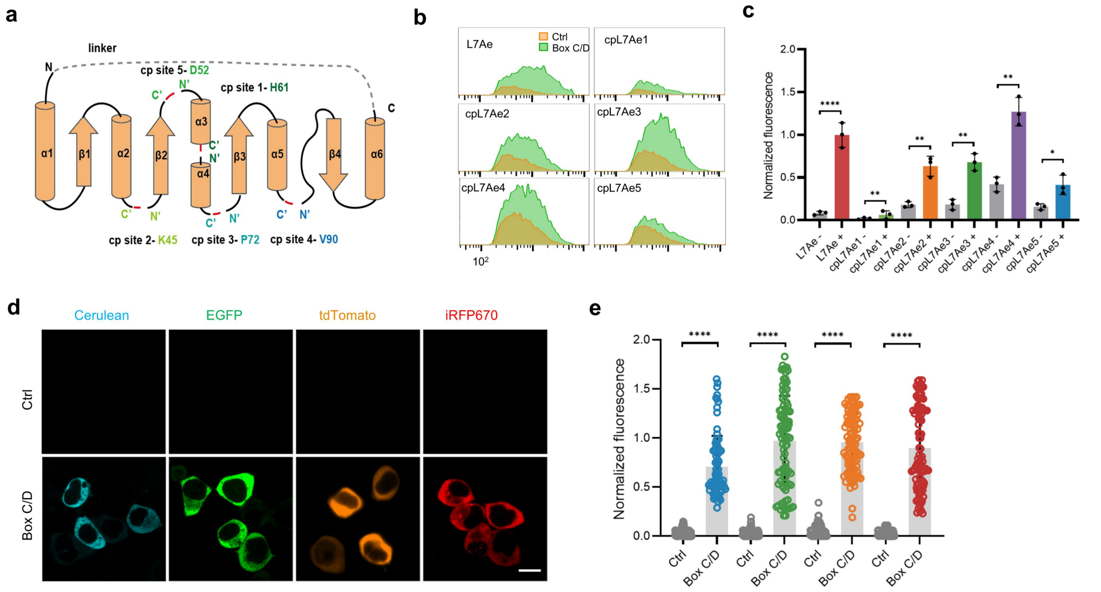


**Supplementary Fig. 15 FLIPS system based on L7Ae and cognate RNA motif. a**, Schematic showing circular permutation strategy to create fL7Ae variants with RNA-binding site in close proximity to the degron. The original N and C termini were connected by a linker, and permutation sites were indicated. **b**, Flow cytometry profiles for HEK293T cells expressing EGFP – L7Ae1-5 – R$_6$RG with circular control RNA (-) or circular Box C/D motif (+). **c**, Normalized EGFP fluorescence intensity in (**b**). Data are normalized to mean fluorescence intensity for cells expressing EGFP–L7Ae–R$_6$RG with circular Box C/D motif. **d**, Confocal images for cells expressing fL7Ae fused with different fluorescent proteins for multicolor imaging. Scale bars, 10 μm. **e**, Normalized fluorescence intensities for individual cells (100 cells from three independent experiments) in (**d**). Data are normalized to mean fluorescence intensity for cells expressing Cerulean – fL7Ae and circular RNA. **c**,**e**, Statistical analysis was performed using a two-tailed t-test (*P < 0.1, **P < 0.01, ****P < 0.0001). Error bars represent s.d. of three independent experiments.

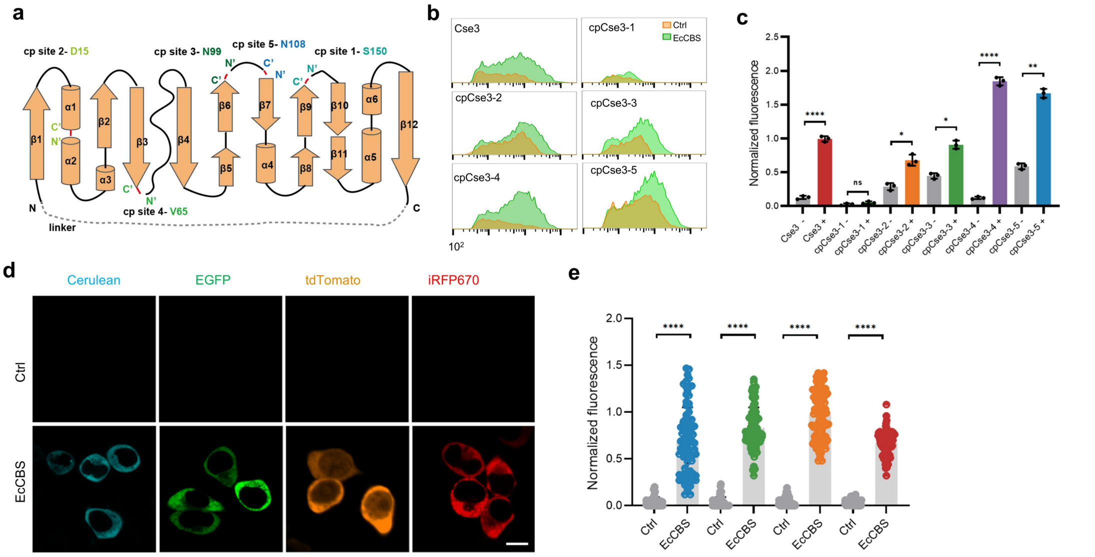

**Supplementary Figure 16. Engineering of FLIPS system based on Cse3 and cognate RNA motif. a**, Schematic showing circular permutation strategy to create fCse3 variants with RNA-binding site in close proximity to the degron. **b**, Flow cytometry profiles for HEK293T cells expressing EGFP – Cse3 variant-$R_6$RG with circular control RNA (-) or circular EcCBS motif (+). **c**, Normalized EGFP fluorescence intensity in (**b**). Data are normalized to mean EGFP fluorescence intensity for cells expressing EGFP–Cse3–$R_6$RG with circular EcCBS motif. **d**, Confocal images for cells expressing fCse3 fused with different fluorescent proteins for multicolor imaging. Scale bars, 10 µm. **e**, Normalized fluorescence intensities for individual cells (100 cells from three independent experiments) in (**d**). Data are normalized to mean fluorescence intensity for cells expressing Cerulean–fCse3 and circular RNA. **c**,**e**, Statistical analysis was performed using a two-tailed t-test (n.s., not significant, *P < 0.1, **P < 0.01, ****P < 0.0001). Error bars represent s.d. of three independent experiments.

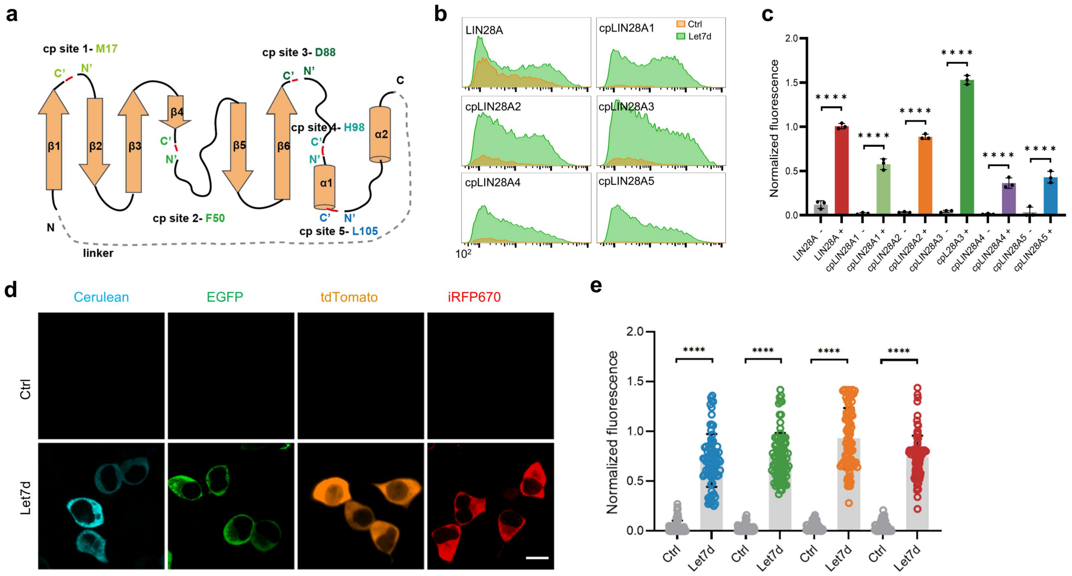


**Supplementary Fig. 17 Engineering of FLIPS system based on LIN28A and cognate RNA motif. a**, Schematic showing circular permutation strategy to create fLIN28A variants with RNA-binding site in close proximity to the degron. **b**, Flow cytometry profiles for HEK293T cells expressing EGFP−LIN28A variant-$R_6$RG with circular control RNA (-) or circular Let7d motif (+). **c**, Normalized EGFP fluorescence intensity in (**b**). Data are normalized to mean EGFP fluorescence intensity for cells expressing EGFP−LIN28A−$R_6$RG with circular Let7d motif. **d**, Confocal images for cells expressing fLIN28A fused with different fluorescent proteins for multi-color imaging. Scale bars, 10 μm. **e**, Normalized fluorescence intensities for individual cells (100 cells from three independent experiments) in (**d**). Data are normalized to mean fluorescence intensity for cells expressing Cerulean−fLIN28A and circular RNA. **c**,**e**, Statistical analysis was performed using a two-tailed t-test (****$P < 0.0001$). Error bars represent s.d. of three independent experiments.

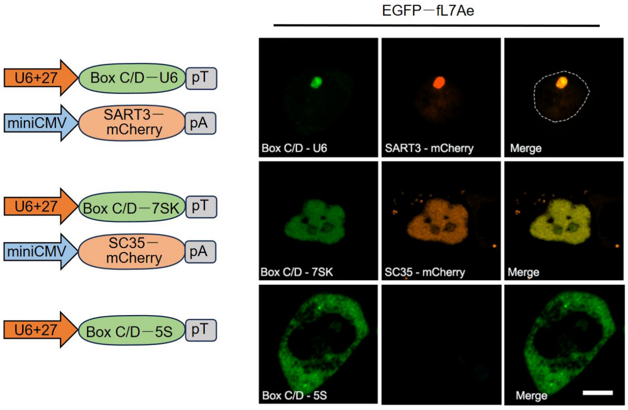


**Supplementary Fig. 18 Effect of fL7Ae system on subcellular localization of RNAs.** Confocal images for cells expressing U6 splicing RNA, 7SK small nuclear RNA or 5S ribosomal RNA and fL7Ae system. The cells were co-transfected with SART3 – mCherry and SC35 – mCherry to label nuclear speckles and Cajal body, respectively. Scale bar, 5 μm. Representative data from three independent experiments.

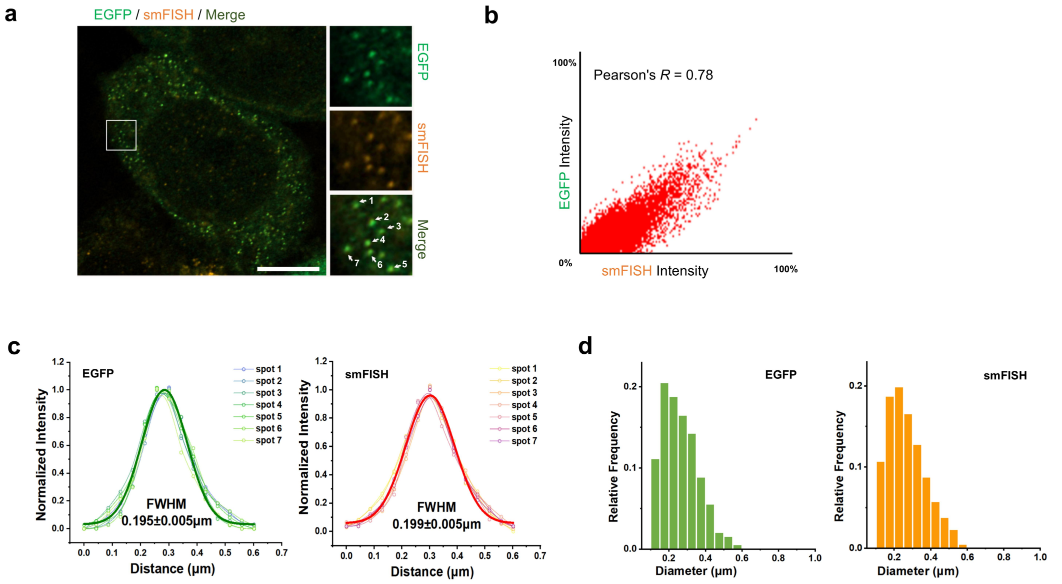


**Supplementary Fig. 19 Single-molecule imaging of iRFP670 mRNA with 24×Box C/D tags using fL7Ae system in comparison to smFISH approach. a**, Confocal images for fixed MCF-7 cells co-expressing fL7Ae system and iRFP670 mRNA with 24×Box C/D tags, and colocalization with TAMRA-labeled smFISH probes specific to the iRFP670 sequence. Zoom of selected region in the white box was shown on the right. Representative 25 cells from three independent experiments. Scale bar, 5 μm. **b**, Scatter plot showing the co-localization of EGFP and smFISH signals in (**a**). **d**, Normalized fluorescence intensity profiles for EGFP and TAMRA puncta marked by white arrows in (**c**). Symbols, experimental data; thin lines, Gaussian fits to the individual fluorescence intensity profiles; thick line, Gaussian fit to the average profiles, yielding a full width at half maximum (FWHM) of 0.195±0.05 μm for EGFP puncta and 0.199±0.05 μm for TAMRA puncta, respectively. **e**, Histogram of EGFP puncta diameter (Left) and TAMRA puncta diameter (Right), N=1800 puncta in 10 cells from three independent experiments.

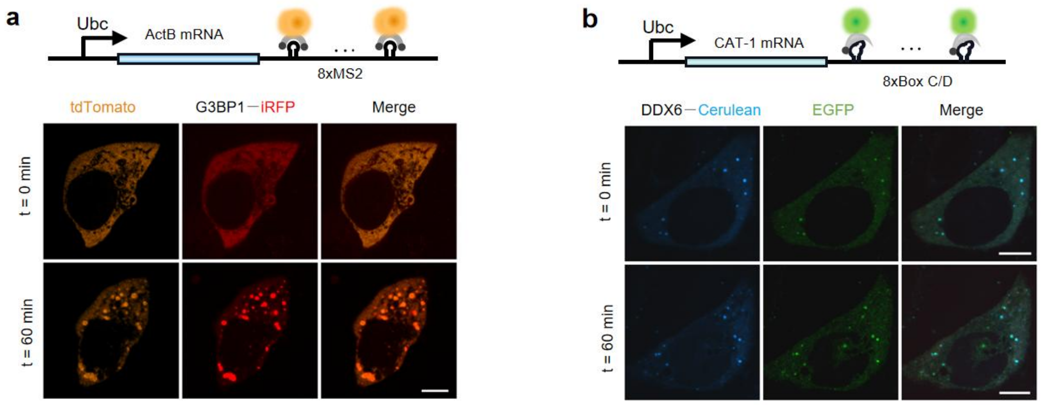


**Supplementary Fig. 20 Two-color imaging of β-actin and CAT-1 translocation upon induction. a**, Plasmid construct for expressing ActB – 8×MS2 mRNA and confocal images for MCF-7 cells co-expressing ActB – 8×MS2, tdTomato-fMCP and the SG indicator G3BP1 – iRFP before and after arsenite (500 μM) induction for 60 min. Scale bar, 5 μm. **b**, Plasmid construct for expressing CAT-1–8× Box C/D mRNA and confocal images for MCF-7 cells expressing CAT-1–8× Box C/D, EGFP–fL7Ae and the PB indicator DDX6 – Cerulean before and after arsenite (500 μM) induction for 60 min. Scale bar, 5 μm.

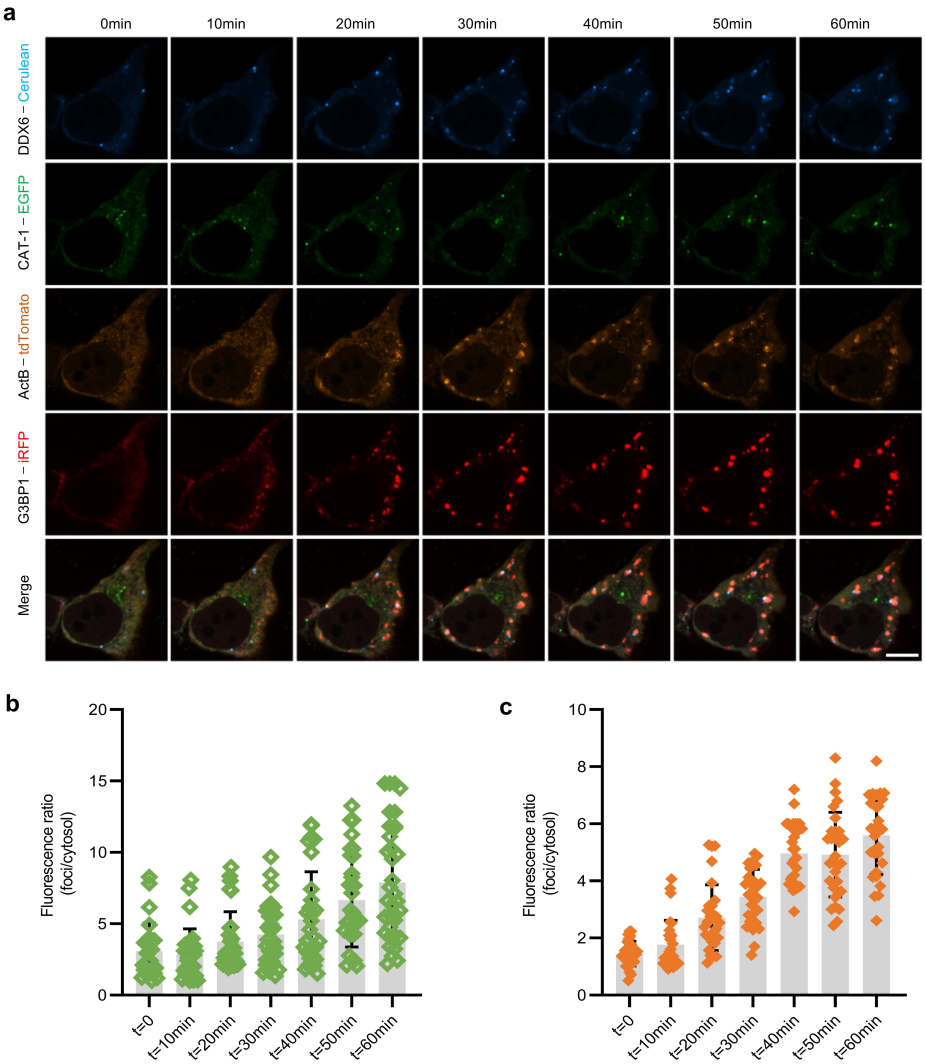


**Supplementary Fig. 21 Real-time imaging of β-actin and CAT-1 dynamic translocation using FLIPS system. a**, Confocal images of cells co-expressing ActB – 8×MS2, tdTomato – fMCP, CAT-1-8×Box C/D and EGFP – fL7Ae at different time intervals upon arsenite (500 μM) induction. Scale bar, 5 μm. **g**, Analysis of ratios of EGFP or tdTomato fluorescence intensities in the foci over those in the cytosol (foci/cytosol) in (**a**). Statistical analysis was performed using a two-tailed t-test (****P<0.0001). Data are represented as mean ± s.d. from three independent experiments.
